# The Hidden Diversity of Vascular Patterns in Flower Heads

**DOI:** 10.1101/2023.10.10.561718

**Authors:** Andrew Owens, Teng Zhang, Philmo Gu, Jeremy Hart, Jarvis Stobbs, Mikolaj Cieslak, Paula Elomaa, Przemyslaw Prusinkiewicz

## Abstract

- Vascular systems are intimately related to the shape and spatial arrangement of the plant organs they support. We investigate the largely unexplored association between spiral phyllotaxis and the vascular system in Asteraceae flowers heads.
- We imaged heads of eight species using synchrotron-based X-ray micro-computed tomography and applied original virtual reality and haptic software to explore head vasculature in three dimensions. We then constructed a computational model to infer a plausible patterning mechanism.
- The vascular system in the head of the model plant *Gerbera hybrida* is qualitatively different from those of *Bellis perennis* and *Helianthus annuus*, characterized previously. *Cirsium vulgare, Craspedia globosa, Echinacea purpurea, Echinops bannaticus*, and *Tanacetum vulgare* represent variants of the Bellis and Helianthus systems. In each species the layout of the main strands is stereotypical, but details vary. The observed vascular patterns can be generated by a common computational model with different parameter values.
- In spite of the observed differences of vascular systems in heads, they may be produced by a conserved mechanism. The diversity and irregularities of vasculature stand in contrast with the relative uniformity and regularity of phyllotactic patterns, confirming that phyllotaxis in heads is not driven by the vasculature.

## Introduction

The development of vascular tissues is one of the key evolutionary innovations that resulted in the explosive radiation of plants on Earth. Vascular tissues, composed of xylem and phloem, provide mechanical support for plants as well as a route for the transport of water, nutrients and signaling molecules such as hormones (Lucas *et al*., 2013). The vascular system extends throughout an entire plant (Esau, 1965a,b). At the level of individual organs, leaf venation patterns have received much attention due to their complexity, diversity, and easy accessibility to direct observation. At the whole-plant level, studies have focused on vasculature in roots and shoot stems.

Until recently, the analysis of volumetric vascular structures involved the laborious process of obtaining serial tissue sections and inferring the three-dimensional structure of the vascular system from them (Kaplan, 1937; Kang *et al*., 2003). Computed tomography alleviates this methodological difficulty by directly providing three-dimensional data stacks (Stuppy *et al*., 2003; Lee *et al*., 2006). In this paper we employ laboratory-based and synchrotron-radiation-based X-ray micro-computed tomography (Withers *et al*., 2021) complemented with computational modeling to investigate the vascular patterns of flower heads. While micro-CT has been found useful for studying internal plant structures (e.g., Dhondt *et al*., 2010; Karunakaran *et al*., 2015; Prunet and Duncan, 2020; Piovesan *et al*., 2021) including stem vasculature (Brodersen *et al*., 2011, Brodersen *et al*., 2012; McElrone *et al*., 2013; Brodersen and Roddy, 2016), the imaging and analysis of flower heads represent a particular challenge due to the geometric complexity of their vascular systems.

Flower heads – the inflorescences in the Asteraceae plant family – typically consist of hundreds of small, densely packed florets attached to an enlarged receptacle and surrounded by protective involucral bracts. The involucral bracts and florets are arranged into intersecting left- and right-winding spirals (parastichies), which frequently occur in consecutive Fibonacci numbers (Jean, 1994; Barabé and Lacroix, 2020). The geometric regularity and mathematical properties of these patterns have attracted multidisciplinary interest for centuries (Adler *et al*., 1997). They have also raised questions regarding the organization of the vascular system that supplies the florets with water and nutrients. Historically first, Philipson (1946) reported that the primary vascular strands in the heads of *Bellis perennis* (common daisy) formed a reticulate network aligned with the parastichies, i.e., with the main vascular strands parallel to the spiral phyllotactic arrangement of the florets. Subsequently, Philipson (1948) reported a similar pattern in two other members of the Aster family: *Hieracium boreale* and *Dahlia gracilis*. In contrast, Durrieu *et al*. (1985) found that the vascular network in the heads of *Helianthus annuus* (common sunflower) was organized sectorially, with the main vascular strands spreading radially from the stem towards the head rim, and the veins from involucral bracts and florets attaching to these strands irrespective of the parastichy pattern. The sectorial organization of the sunflower head vasculature was confirmed by Alkio *et al*. (2002), who demonstrated that photoassimilates tagged with radioactive carbon propagated to achenes (seeds) along radial sectors, as opposed to following the parastichy pattern.

The discrepancy between the vasculature of Bellis and sunflower heads raised our interest in the type of vasculature present in *Gerbera hybrida*, a model plant used to study the phyllotaxis of heads (Zhang *et al*., 2021). Using synchrotron-radiation based X-ray micro-computed tomography (SR-μCT) imaging, we show that gerbera head vasculature is also organized sectorially. Unexpectedly, however, this organization is qualitatively different from that of the sunflower heads reported by Durrieu. To exclude the possibility that the observed distinction is an artefact of different experimental methods, we also imaged sunflower and Bellis heads. Our results confirm those reported previously while augmenting Philipson’s description of Bellis with a more detailed observation of the position of the vascular strands with respect to the florets. Intrigued by the differences between gerbera, sunflower and Bellis, we have additionally imaged the heads of five previously unstudied species: *Cirsium vulgare* (common thistle), *Craspedia globosa, Echinacea purpurea* (coneflower), *Echinops bannaticus, and Tanacetum vulgare*. We show that their vascular systems are variants of, or intermediate between, those observed in Bellis and sunflower.

The complexity and diversity of vascular patterns in heads lead to the question of how these patterns develop. Data (Zhang *et al*., 2021) suggest that the basic patterning processes in heads are similar to those found in the inflorescences of plants not forming heads, such as Arabidopsis (Reinhardt *et al*., 2003), tomato (Bayer *et al*., 2009) and *Brachypodium* (O’Connor *et al*., 2014). The key patterning mechanism is the interaction between the plant hormone auxin and its cellular transporters, primarily the PIN1 efflux carriers. It leads to the emergence of auxin concentration maxima that organize the incipient primordia into a phyllotactic pattern, and canals of auxin transport that connect these primordia to the already formed vasculature, patterning new strands. These processes are best understood with the help of computational models (Prusinkiewicz and Runions, 2012). However, the existing models that simulate vascular patterns directly at the level of molecular interactions only capture relatively simple patterns and pattern elements, such as canals connecting auxin sources to sinks, and operate on two-dimensional leaf blades or plant sections (Runions *et al*., 2014). To overcome this limitation and capture the complex three-dimensional vascular architecture of flower heads, we have constructed a higher-level model in which molecular interactions are distilled into geometric rules. Our model integrates and extends the different classes of models proposed previously (Cieslak *et al*., 2021), most significantly the resistance-based model of leaf venation introduced by Runions *et al*. (2017). It reproduces the patterns found in the observed heads, provides a plausible high-level explanation of pattern self-organization during head development, and shows that the diversity of the observed patterns may result from relatively small variations of a common patterning mechanism.

## Materials and Methods

### Plant material

*Gerbera hybrida* cv. ‘Terra Regina’, the transgenic gerbera lines expressing the constructs for the antisense-*GRCD2* line TR15 (Uimari *et al*., 2004; Zhang *et al*., 2017), the *DR5rev:3xVENUS-N7* line TR3 (Zhang *et al*., 2021), as well as *Helianthus annuus* cv. ‘Pacino cola’ and *Craspedia globosa* plants were grown under standard greenhouse conditions. Samples from *Bellis perennis, Echinacea purpurea* cv. ‘White Swan’, *Echinops bannaticus, Tanacetum vulgare* and *Cirsium vulgare* plants were collected from outdoor grown plants in Helsinki, Finland. For each species, fully patterned flower heads were collected in two biological replicates. Additionally, developmental series of head samples were collected for sunflower and Bellis. The smallest samples with embedded early-stage head meristems (diameter < 1 mm) were randomly dissected from the plants under a stereomicroscope, and the developmental stage of the meristems was estimated on the basis of the micro-CT imaging. Following the collection, the samples were fixed, dehydrated, critical-point dried and mounted on scanning electron microscopy (SEM) stubs as described by Zhang *et al*. (2021).

### Microcomputed tomography

Head samples were scanned using laboratory-based or SR-based X-ray micro-CT. The choice of the technique was affected by its availability and technological progress over the duration of our study (2016 to 2022), with a preference for SR-μCT due to the superior quality of the resulting images. The laboratory-based scanners were a Skyscan 1272 micro-CT scanner (Bruker Micro-CT, MA, USA) and a GE Phoenix Nanotom scanner (GE Sensing & Inspection Technologies GmbH, Germany). Synchrotron scanning was conducted at the Biomedical Imaging and Therapy (BMIT) facility at the Canadian Light Source, Saskatoon, Canada, https://bmit.lightsource.ca. We used the bending magnet beamline (05B1-1) with the X-ray energy of approximately 20kV, which had been found optimal for imaging plant samples (Kanurakaran *et al*., 2015). The beamline produces a monochromatic and partially coherent beam, which we employed to enhance contrast using phase-contrast imaging. Details concerning the synchrotron scanning are given in the Supplementary Text S1, and attributes pertinent to the scanning and visualization of individual heads are listed in Table S1.

### Tomographic image reconstruction

Three-dimensional image stacks were reconstructed from the projection images using NRecon version 1.7.0.1 (Bruker Micro-CT, MA, USA) or the Tofu image processing toolkit (Farago *et al*., 2022) included in the UFO-KIT 0.9 package (Vogelgesang *et al*., 2016) augmented with the EZ-UFO graphical interface (https://github.com/sgasilov/ez_ufo, S. Gazilov, Canadian Light Source), using default configuration parameters. The methods applied to the specific samples are indicated in Table S1. Ring artifacts in the tomograms were suppressed using a low-pass Gaussian filter. Multiple sections of tall samples were combined into one image stack using ImageJ/Fiji (Schindelin *et al*., 2012), by visually matching scans in the overlapping portions of each section. To reduce file sizes, each image stack was cropped to eliminate excessive background and reduced from 32-bit to 16-bit representation by selecting the relevant portion of the histogram using ImageJ/Fiji. The final stacks were assembled into tiff (Tag Image File Format) files. The reconstructions were carried out using Dell Alienware Area-51 Desktop PC with a 3.1 GHz Intel Core i9-7940X CPU, 64 GB RAM, and a NVIDIA GeForce RTX 2080 Ti graphics card with 11 GB VRAM, running the Windows 10 Enterprise operating system (NRecon reconstructions) or Linux Ubuntu 18.04 (UFO-KIT reconstructions).

### 3D Image analysis, segmentation and visualization

The reconstructed image stacks were analyzed using ImageJ/Fiji, MorphographX (Barbier de Reuille *et al*., 2015), and two experimental programs, SHVR (Gu, 2022) and ViNE (Hart, 2020), developed at the University of Calgary. SHVR operates directly on volumetric data and facilitates segmentation of vascular strands by using a haptic device (3D Systems Touch, Rock Hill, SC) to give the operator the impression of physically touching and following them. ViNE operates on polygonal meshes extracted from volumetric data and facilitates segmentation by using a virtual reality system, which gives the operator the impression of being immersed in the flower head and segmenting vascular strands directly in 3D. The isosurface meshes were generated using a custom-made IsoPoly program utilizing the OpenVDB library (https://www.openvdb.org). The final images were rendered directly by SHVR or ViNE, or were output as PLY (Polygon File Format) files and rendered using Blender v2.8 (https://blender.org). Movies were assembled using ffmpeg (https://www.ffmpeg.org). While diverse hardware was used for software development and research, the results have been tested and are reproducible on the Dell Alienware desktop PC running Windows 10 Enterprise (as specified above) coupled with the 3D Systems (Rock Hill, SC) Touch haptic device (for SHVR) or the HTC (New Taipei Corporation, Taiwan) Vive virtual reality system (for ViNE).

### Histological analysis

To obtain additional information of the early stages of head development, we complemented tomographic imaging with histological analysis. Five rosette sectors containing representative head meristems were collected from the transgenic gerbera plants expressing the *DR5rev:3xVENUS-N7* auxin reporter construct (Zhang *et al*., 2021). To visualize the DR5 signals, the samples were first imaged with a Leica FCA M205 fluorescence stereomicroscope (Leica Microsystems) using a GFP filter. The same samples were then fixed, dehydrated, and embedded in paraffin blocks as described by Laitinen *et al*. (2006). Serial 10 μm sections were cut in a horizontal direction using a microtome. The paraffin strips were attached to imaging slides, deparaffinized and stained in 0.2% toluidine blue for 10 mins. After washing with sterile H_2_O for three times, the slides were mounted with 50% glycerol and imaged with a Leica FCA M205 microscope.

### Computational modeling

To investigate the mechanism of vascular patterning in flower heads and the causes of their diversity, we constructed parametrized simulation models of vascular patterning in gerbera, sunflower and Bellis. Vascular patterning was simulated on data-driven models of growing heads with simulated phyllotactic patterns (Fig. S10). For gerbera we used the flower head model of Zhang *et al*. (2021); for sunflower and Bellis we recalibrated that model using image data obtained by scanning the respective heads at different developmental stages. All models were written in C++ extended with constructs of the L+C plant modeling language (Karwowski and Prusinkiewicz, 2003; Prusinkiewicz *et al*., 2007), and executed using the lpfg simulator incorporated into the Virtual Laboratory (vlab) v5.0 plant modeling environment (http://algorithmicbotany.org, https://github.com/AlgorithmicBotany/vlab). Simulations were performed on MacBook Pro computers under macOS High Sierra. Movies were assembled using ffmpeg.

## Results

### Gerbera flower head displays a previously undescribed sectorial vascular system

A gerbera (cv. ‘Terra Regina’) head has approximately 600-700 individual florets surrounded by 80-90 involucral bracts (Zhang *et al*. 2021) (Fig. 1a,b; for all morphological features see also Movie 1). The bracts and florets form on a mushroom-shaped receptacle with a convex but relatively flat upper (adaxial) surface that meets the lower (abaxial) surface along a discernible rim (Fig. 1c, dashed white line). SR-μCT imaging revealed that the stem supporting the head has approximately 30 longitudinal vascular strands (Fig. S1d,e; Fig. S6e). Upon entering the receptacle, these strands branch and spread radially towards the receptacle rim (Fig. 1d; Fig. S1d,e), and occasionally merge. The outermost strands enter the involucral bracts, where they split into parallel veins running the length of each bract. The intermediate strands, enter ray and trans florets near the head rim. The innermost strands change direction and extend towards the receptacle center, forming a system of adaxial strands (Fig. 1c; Fig. 2b,c; Fig. S1b,c). Approaching the head center, some of these strands terminate early or merge, such that their density remains approximately constant (Fig. 2b). Individual disc florets connect with the adaxial strands via one or several floret veins, which are approximately perpendicular to the upper surface of the receptacle (Fig. 2b,c). Except close to the rim, the ground tissue (parenchyma) of the receptacle does not contain discernible vasculature (Fig. 1c).

**Fig. 1.**
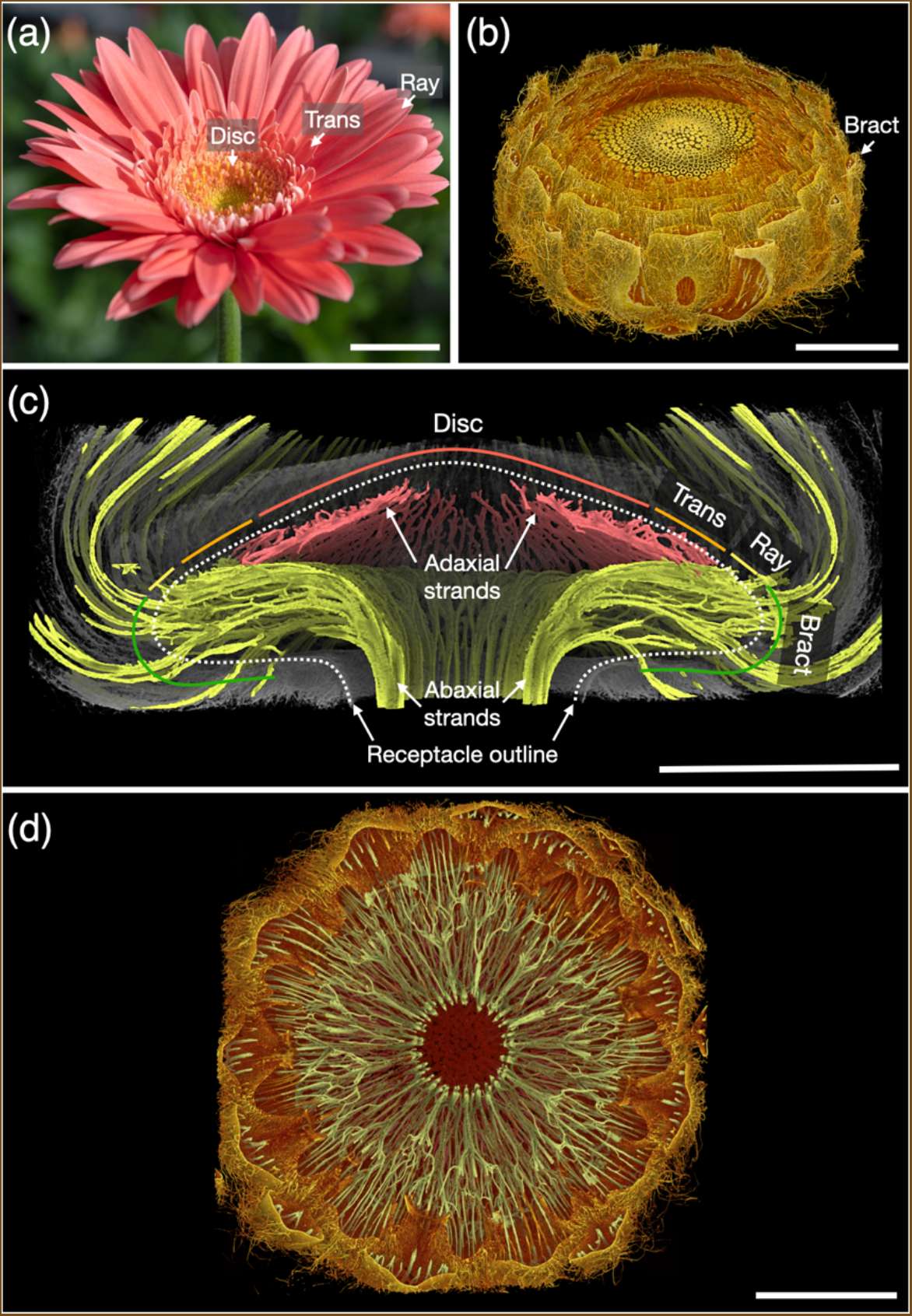
Vascular architecture of gerbera heads. (a) A mature head with three different types of florets (ray, trans, disc) emerging acropetally from the margin of the head. (b) 3D reconstruction of an earlier developmental stage of a gerbera flower head. (c) Longitudinal side view of half of the head with the vascular system segmented using SHVR. The abaxial and adaxial vascular strands are colored green and red, respectively. (d) A virtual transverse cut showing the abaxial vascular system within a gerbera head. The branched vascular system spreads from the stem radially and enters the involucral bracts at the head rim. Scale bars: 2 cm in (a); 5 mm in (b-d).

**Fig. 2.**
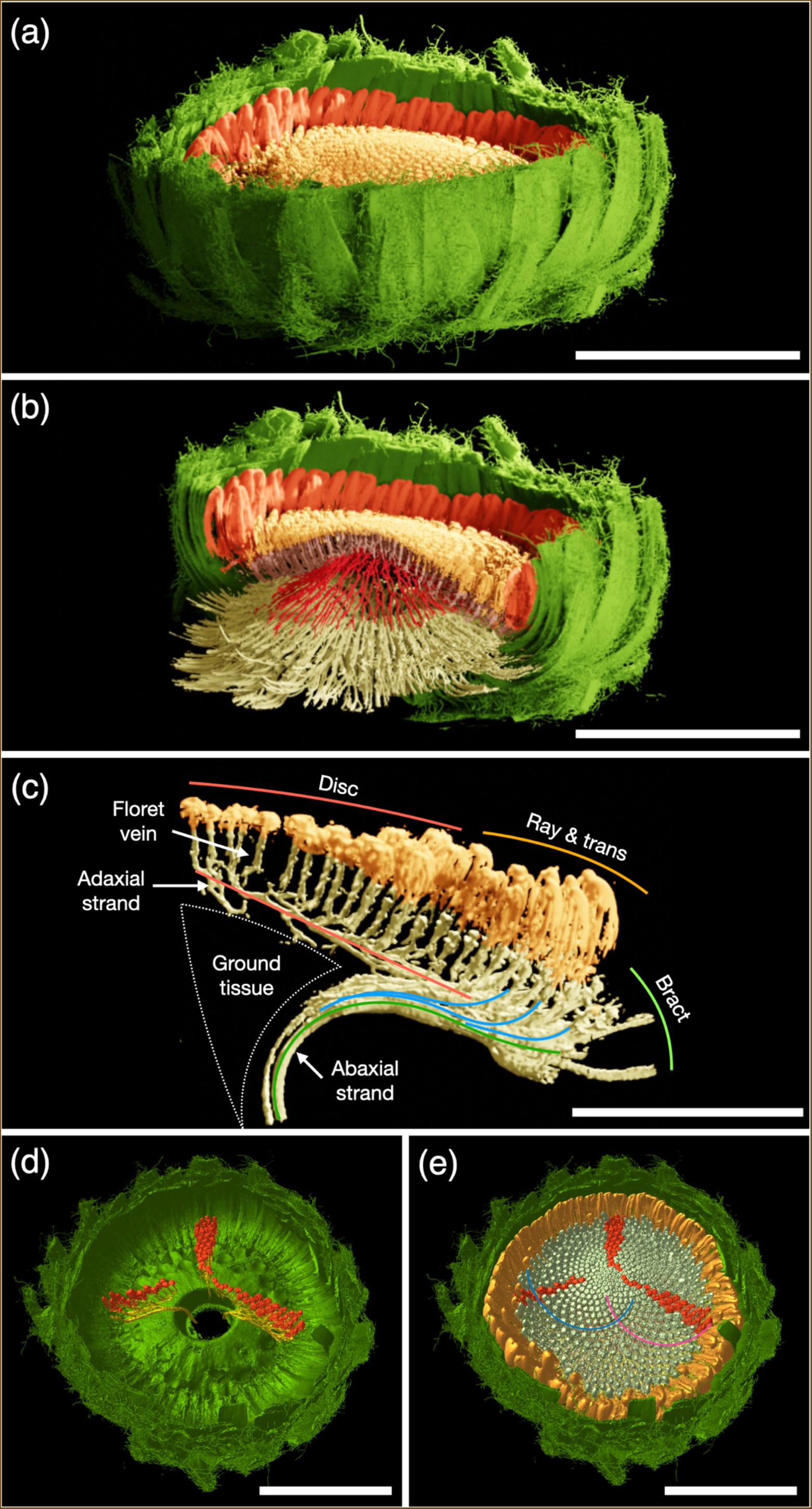
Phyllotaxis and the vascular system in a gerbera flower head. (a) 3D rendering of a scanned gerbera head segmented using ViNE, indicating the bracts (green), ray florets (orange), and disc florets (yellow). (b) A virtual cut exposing the vascular structure of the same head. Abaxial strands (straw-colored) extend outward to the bracts and connect near the head perimeter to adaxial strands (red), which extend towards the head center. Short floret veins (purple), perpendicular to the receptacle surface, connect florets to the adaxial veins. (c) An isolated sector of the vascular system showing adaxial strands (highlighted in red) originating from a single abaxial strand, and the connected floret veins. The abaxial strands connect to bracts (green), and ray and trans florets (blue). (d) A flowerhead from above with all but three vascular strands emerging from the stem removed. The vascular system originating from each strand (yellow) supports a separate sector of florets on the head (red). (e) The same flower head as (d) with all florets displayed, showing that the florets in each sector are from different parastichies. A representative pair of spiral parastichies are highlighted with blue and pink lines. Scale bars: 1 cm in (a,b,d,e); 5 mm in (c).

We further analyzed the connections between vascular strands and florets by segmenting florets connected to selected adaxial strands using the ViNE software (Fig. 2d,e). Our analysis confirmed that the vascular system of the gerbera heads is organized sectorially, as implied by the radial organization of the main abaxial and adaxial strands (Fig.1c,d; Fig. 2b,c). This organization is independent of the spiral phyllotaxis of the head: an individual adaxial strand supplies floret primordia that belong to different parastichies, while florets that belong to different segments of the same parastichy are supplied by different adaxial strands (Fig. 2d,e).

### 3D visualization confirms previously described vascular patterns in Bellis and sunflower

Our analysis indicated that the vascular network in gerbera is qualitatively different from those previously reported in Bellis (Philipson, 1946) and sunflower (Durrieu *et al*., 1985). To confirm that the observed differences are not an artefact of distinct experimental methods, we have imaged, segmented and analyzed Bellis and sunflower heads using SR-μCT as well (Fig. 3). As in gerbera, the florets in these species are arranged into spiral phyllotactic patterns (Fig. 3a,e), although the receptacles have different overall shapes. The receptacle in sunflower (cv. ‘Pacino cola’) is concave with a shallow, slightly depressed central region, which raises toward the peripheral rim (Fig. 3c). By contrast, the receptacle of Bellis is a vertically elongated dome (Fig. 3g).

**Fig. 3.**
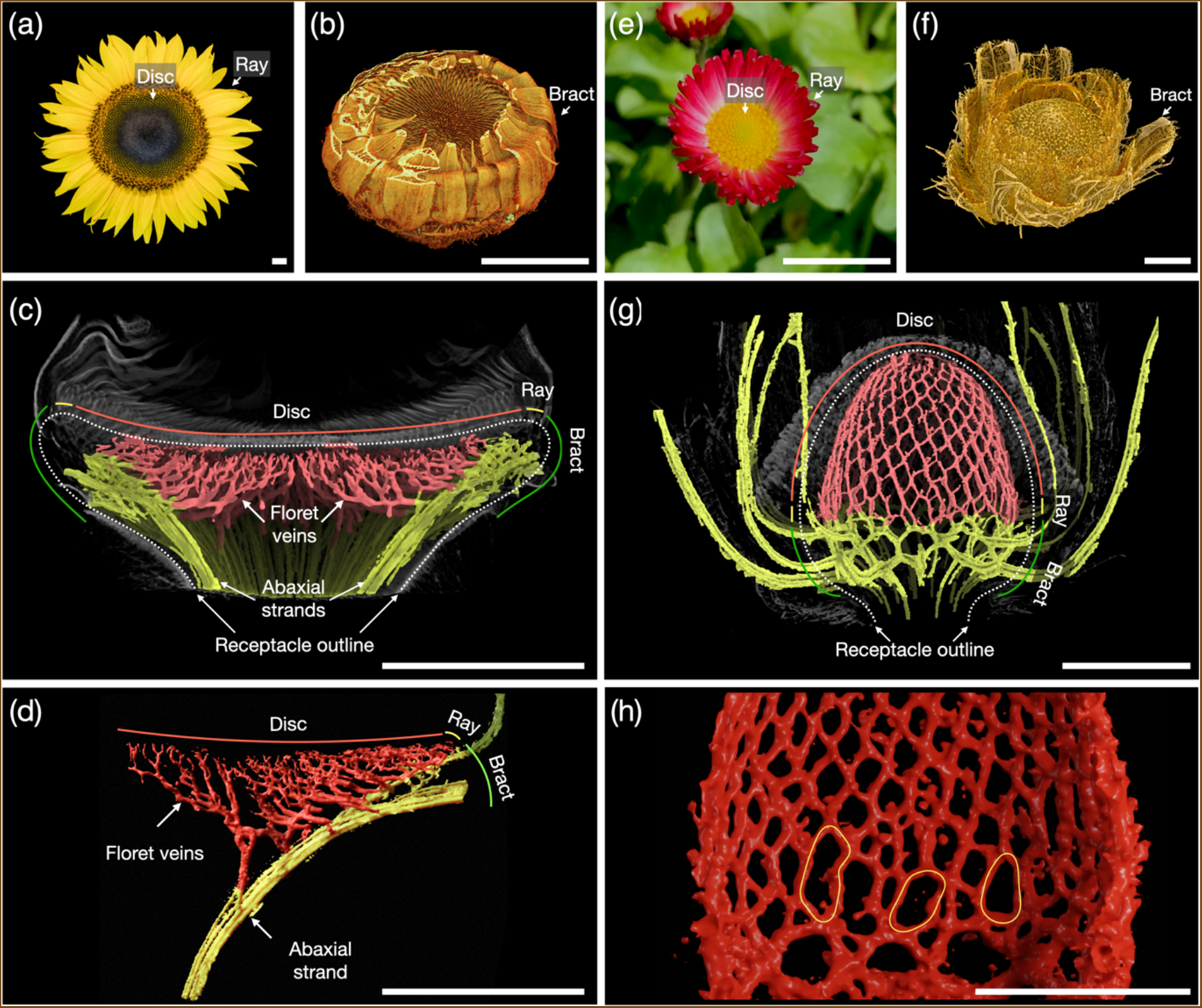
Vascular systems of sunflower (a-d) and Bellis (e-h) heads. (a) A mature flower head of sunflower bearing two different types of florets: ray and disc. (b) 3D-reconstruction of an earlier developmental stage of a sunflower head. (c) Longitudinal view of half of a sunflower head with the vascular system segmented with SHVR. (d) ViNE segmentation showing floret veins connecting to a single abaxial strand in the sunflower head. (e-g) Images of the Bellis head corresponding to (a-c). (h) A magnified view of the reticulate Bellis head venation, highlighting gaps near the base of the vascular system (yellow lines). Scale bars: 1 cm in (a-c,e); 5 mm in (d); and 2 mm in (f,g,h).

Our observations were consistent with those reported previously. As in gerbera, the vasculature of sunflower heads is organized sectorially, with no correspondence between abaxial strands and the phyllotactic pattern (Fig. S2b,c). Large abaxial strands spread outward from the stem to the involucral bracts and peripheral ray florets (Fig. 3c,d; Fig. S2a,b) but, in contrast to gerbera, there are no adaxial strands extending inward: the floret veins connect directly to the abaxial strands across the parenchyma (Fig. 3d). As observed by Durrieu *et al*. (1985), the course of these veins is highly irregular and they often merge, forming a branched system (Fig. 3d).

Likewise, our observations of the vasculature of Bellis generally agree with those of Philipson (1946). The Bellis head vasculature forms a regular reticulate network of vascular strands aligned with the parastichies (Fig. 3g; Fig. S3a,b). However, using SR-μCT we observed more details. Individual floret veins attach to this network near, but not always exactly at, the point of intersection between opposite parastichies (Fig. S3c,d). This regularity breaks near the base of the head, where strands from the stem enter the receptacle and form a less reticulated network. Connections corresponding to one of the two intersecting parastichies are often missing, producing gaps at the base of the network (Fig. 3h, yellow lines).

In summary, the vascular system in gerbera heads exhibits a previously unobserved type of architecture. In addition, our data confirm the previously reported qualitative differences between the vascular systems of Bellis (Philipson, 1946) and sunflower (Durrieu *et al*., 1985) and provide additional insights regarding their three-dimensional structure.

### Vascular patterns in other observed heads are intermediate between sunflower and Bellis

Having observed that flower heads of the three analyzed species have qualitatively different vascular networks, we inquired whether this diversity extends to other Asteraceae species. We analyzed five additional species, resulting in a total of eight species representing two distinct subfamilies of Asteraceae: Cichorioideae (gerbera, thistle and Echinops) and Asteroideae (sunflower, Belli*s*, coneflower, Tanacetum, and Craspedia) (Fig. 4, Fig. S4). None of the additionally scanned species had a vascular system identical to any of the original three species. However, each system could be interpreted as a combination of features from sunflower and Bellis.

**Fig. 4.**
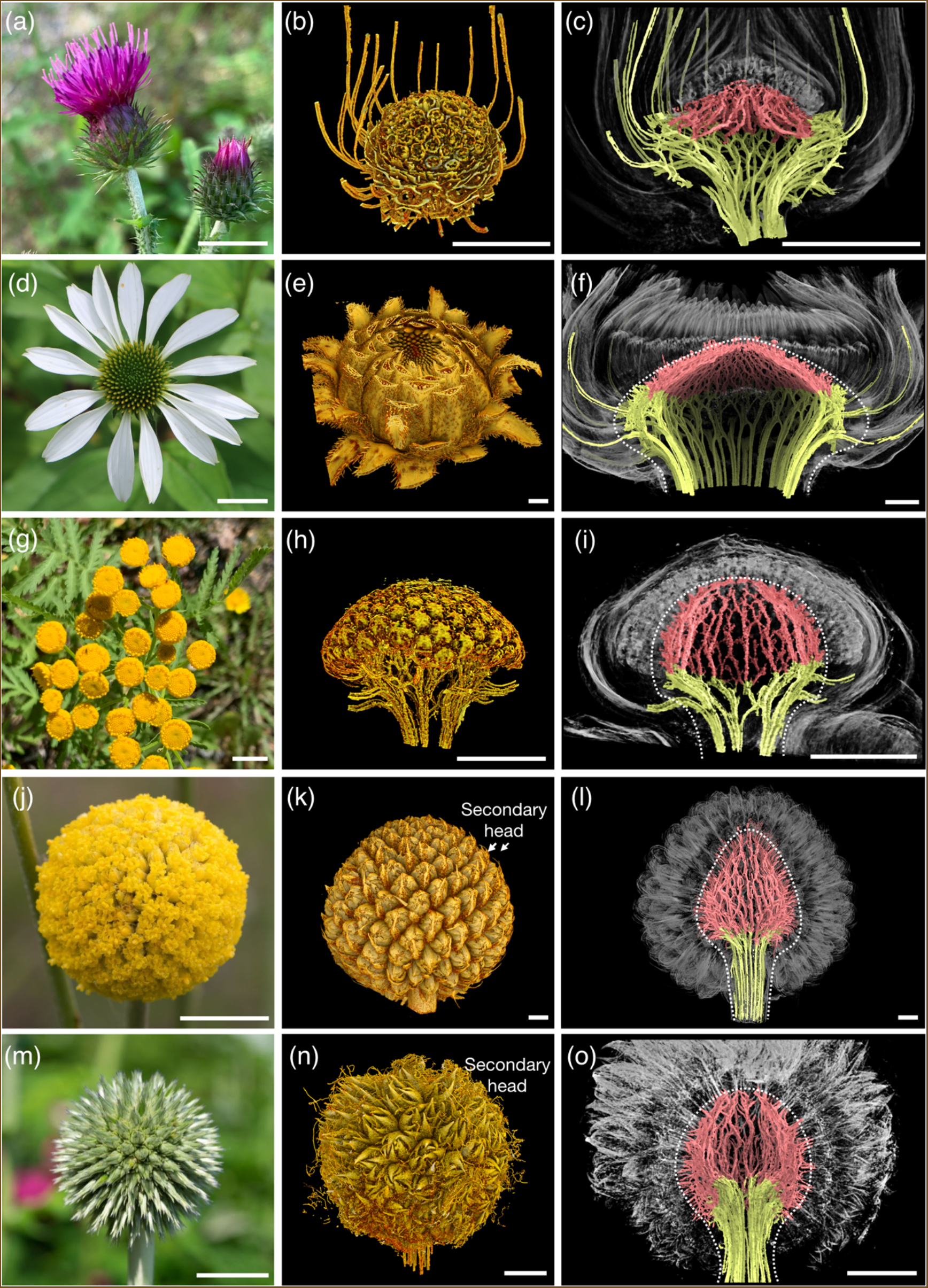
Vascular architecture in: (a-c) thistle, (d-f) coneflower, (g-i) Tanacetum, (j-l) Craspedia and (m-o) Echinops. The first two columns represent mature inflorescences and 3D reconstructions of flower heads that are fully patterned with primordia. The third column shows longitudinal views of half of the respective flower heads with the vascular system segmented using SHVR. In the case of Craspedia and Echinops, the primary heads support secondary heads rather than individual primordia. Scale bars: 1 cm in the first column; 1 mm in the second and third columns.

The receptacle of thistle heads (Fig. 4a-c) is conical. Similar to the sunflower, the floret veins connect directly to the abaxial stem strands (Fig. 4c) and are organized sectorially (Fig. S4a,b).The vascular strands entering the head from the stem extend along the bristly abaxial side of the receptacle and occasionally interconnect (Fig. 4c). These interconnections relate thistle vasculature to Bellis.

Coneflower (Fig. 4d-f) also develops a conical receptacle (Bremer, 1994) and, similar to thistle, has floret veins organized sectorially (Fig. 4f; Fig S4c, top). In contrast to sunflower and thistle, these veins do not self-organize into highly branched structures deeply penetrating the parenchyma (Fig. 3d), but form meandering strands that extend from the head periphery towards the center while remaining close to the adaxial surface of the receptacle (Fig. 4f). The overall course of these strands resembles the adaxial strands in gerbera, except that the gerbera strands are distinct from the floret veins in a manner implying monopodial branching, whereas in coneflower this distinction is absent, implying sympodial branching. Abaxial veins in the coneflower form a reticulate network (Fig S4c, bottom) similar to that found near the base of Bellis heads.

The receptacle of Tanacetum (Fig. 4g) is more elongated (Fig. 4h,i) than that of coneflower or thistle. The vasculature combines features of a sectorial system, with thick strands running approximately in the radial direction, and reticulation manifest in interconnections between these strands (Fig. 4i; Fig. S4d). In contrast to thistle and coneflower, these interconnections appear not only in the abaxial strands, but also in the adaxial strands. In contrast to Bellis, the resulting network is not related to the parastichies.

We also investigated two species with globular receptacles and heads representing syncephalia, i.e., compound structures in which entire heads develop within a head (Zhang and Elomaa, 2021). In Craspedia (Fig. 4j-l), the primary receptacle supports secondary heads with 8 to 10 florets (Bremer, 1994), whereas in Echinops (Fig. 4m-o) the primary receptacle supports second-order heads composed of a single floret subtended by a bract. In both cases, the vascular strands from the stem attach directly to the marginal secondary heads, as the first-order heads lack involucral bracts. The (primary) vascular system of Craspedia is reticulated in a manner resembling Bellis, although it is less regular and vertically more stretched (Fig. S4e). Due to a relatively small number of secondary heads, the vascular system in Echinops is simpler, with the main strands meandering as in Tanacetum and occasionally interconnecting as in Craspedia (Fig. S4f).

### The ontogeny of the vascular system correlates with receptacle growth

To understand how the vascular systems arise, we followed the head development of gerbera, sunflower and Bellis over time (Fig. 5). At an early developmental stage, right after the reproductive transition, the receptacles in all three species are dome-shaped. The stem strands are already visible and connect directly to the early bract primordia (Fig. 5a,d,g).

**Fig. 5.**
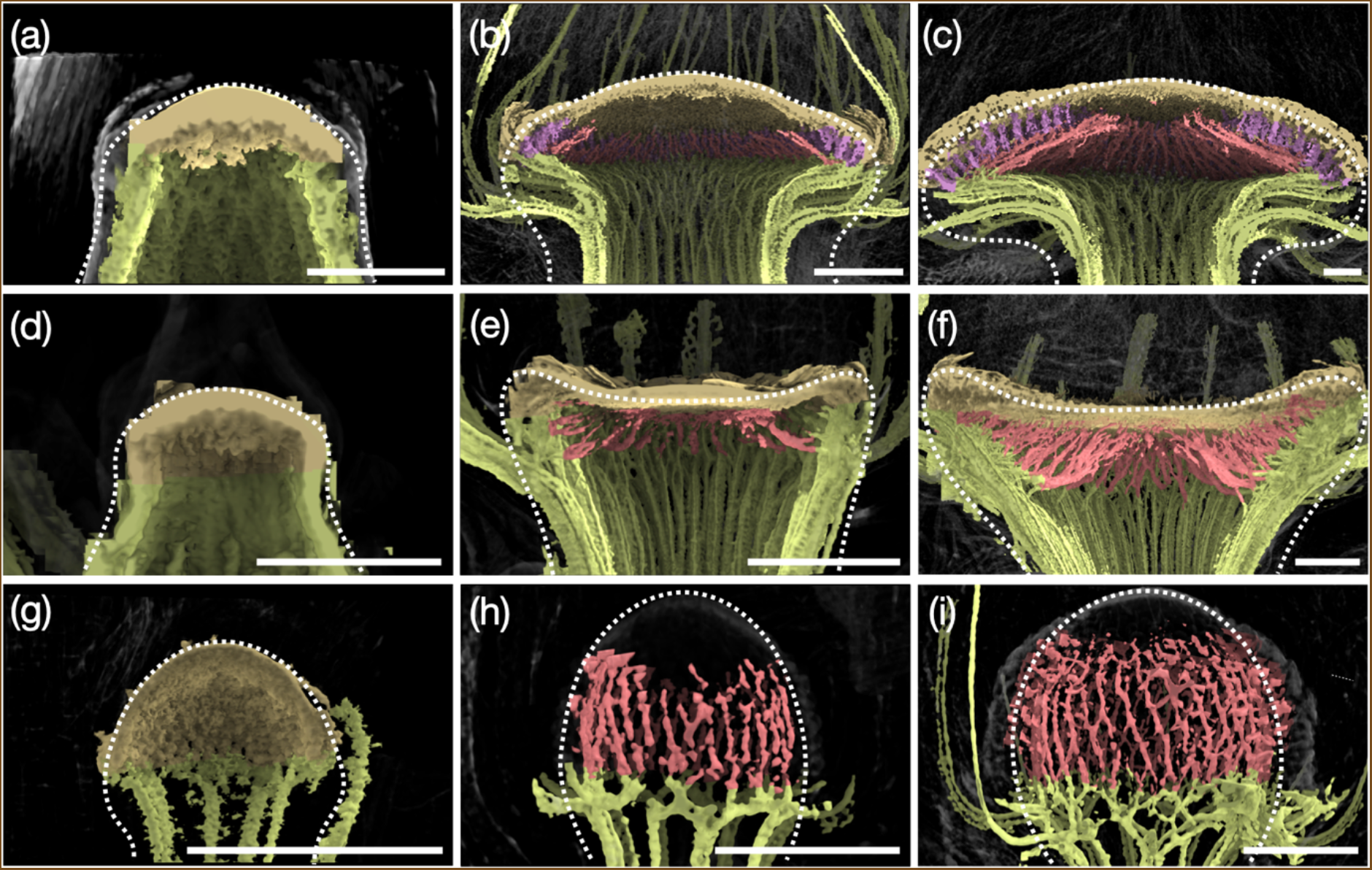
Ontogeny of the vascular systems at different stages of head development in gerbera (a-c), sunflower (d-f) and *Bellis* (g-i). The receptacle outlines are marked in dashed white lines. Scale bars: 0.5 mm.

As development continues, both the florets that gradually fill the meristem surface, and the vascular system that supports them progress from the periphery to the receptacle center (Fig. 5b,e,h). At this stage, the development in the three head types diverge. The gerbera receptacle gradually assumes a convex mushroom shape. This transformation spreads the distal parts of the stem strands outward, producing radially-oriented abaxial strands. The tips of the innermost strands turn toward the center of the receptacle, producing adaxial strands (Fig. 5b,c). Short floret veins then connect the emerging florets to the adaxial strands (Fig. 5c). To examine this process further, we analyzed the scans of a transgenic gerbera line, in which the centre of the inflorescence meristem remains undifferentiated and does not initiate new floret primordia due to downregulation of *SEPALLATA*-like *GRCD* genes (Uimari *et al*., 2004; Zhang *et al*., 2017; Fig. S5). Both the abaxial stem veins and the adaxial veins in these transgenic plants develop as in wild-type gerbera. However, in the head centre, the adaxial veins continue to develop towards the centre despite of the absence of the floret primordia, and eventually they merge below the undifferentiated meristematic surface closing the network (Fig. S5b,c). Altogether, our data suggests that the centripetal veins in gerbera act as a leading strand, to which the veins originating in the florets eventually connect.

In contrast to gerbera, the sunflower heads gradually assume a concave shape (Fig. 5). As florets are patterned in progression towards the center, they become directly connected to the large vascular strands from the stem (Fig. 5e,f).

Vascular strands in the globular head of Bellis form a reticulate structure aligned with two intersecting families of parastichies. During development, the strands aligned with one family become visible first (Fig. 5h), and the intersecting strands following the opposite family emerge later (Fig, 5i), eventually producing the reticulate pattern (Fig. 3g).

### Phyllotactic patterning precedes vascular development in gerbera

Volumetric imaging reveals that organ primordia and vascular strands are already present in the early developmental stages of head development (Fig. 5a,d,g, Fig. S6). To further explore the association between vascular patterning and phyllotaxis, we examined a transgenic gerbera line expressing the *DR5rev:3xVENUS-N7* auxin reporter (Zhang *et al*., 2021). Using this line, we first identified the number of auxin maxima on the meristem surface, then performed serial sectioning and histological staining of the same meristem samples (Figs. S7 and S8). In the earliest developmental stage, the first three DR5 maxima (corresponding to future bract primordia) were patterned (Fig. S7a,e). Histological analysis of the sample showed a continuous ring of densely stained cells at the base of the meristem (Fig. S7i). This region represents the procambial or meristematic ring initiating new vascular strands (Esau, 1954). Subsequently new DR5 maxima emerged, while the procambial region became separated into discrete vascular strands (Fig. S7b-d,f-g,j-l). Their number increased as the head grew, indicating that the new strands were inserted intercalarily between the existing ones (Fig. S7i-l). The number of differentiated vascular strands was always smaller than the number of auxin maxima on the meristem surface, demonstrating that auxin patterning on the surface occurs prior to the differentiation of vascular strands beneath it. A re-analysis of the live-imaging data of gerbera head meristem from Zhang *et al*. (2021) has further shown that the DR5 signals emerge first on the meristem surface, then gradually extend towards the inner tissues (Fig. S9). Taken together, these observations indicate that the specification of vasculature in gerbera, and likely also in other flower heads, shares a conserved auxin-driven patterning mechanism with other species (Bayer *et al*. 2009; O’Connor *et al*., 2014).

### A common model captures vascular pattern development in diverse flower heads

The complexity and diversity of vascular patterns in flower heads lead to the question of the underlying developmental mechanism. To identify a plausible candidate, we explored several hypotheses using a computational model. The model operates on a dynamic template of receptacle growth, simulated using previous data for gerbera (Zhang *et al*., 2021) and new data for sunflower and Bellis (Fig. S10). The input data also include simulated dynamic patterns of primordia emergence. For gerbera it is the pattern described by Zhang *et al*. (2021) (Fig. S10c). For the sunflower and Bellis, the phyllotactic patterns have been simulated using the same algorithm, with parameters modified to capture the characteristics of each species (Fig. S10f,i).

Consistent with the observations (Fig. 5), simulations of vascular pattern development proceed through two phases. In the first phase, the emerging vascular strands connect the early primordia (which may develop into involucral bracts) to the procambial ring positioned at the base of the young head. In the second phase, vascular strands originating from the floret primordia veins connect to the previously formed strands. As the head grows, all strands elongate and deform to accommodate the changes in size and shape of the growing receptacle.

We began with the highly regular pattern observed in Bellis heads. The pattern consists of two families of vascular strands aligned with left- and right-winding parastichies (Fig. 3g). These families are not equivalent: the steeper strands are more pronounced and develop before the strands following parastichies running in the opposite direction (Fig. 5h,i). The question thus arises, on what basis the patterning process distinguishes these strands.

The simplest hypothesis is that each incipient primordium connects first to the closest point on a previously formed strand. With primordia arranged into a spiral phyllotactic pattern, this point coincides with, or lies near, the closest preceding primordium. However, as the distance of the incipient primordium to its closest neighbors along either contact parastichy is approximately the same, the resulting algorithm produces irregular strands, which reflect inconsistent choices of the parastichy to follow (Fig. S11a).

Seeking an alternative, we hypothesized that each new connection minimizes the resistance to auxin transport from the auxin source located at the incipient primordium to the sink located at the receptacle base. A similar model has been proposed for leaf vasculature (Runions *et al*., 2017). The paths of emerging connections then depend not only on distances, but also on the resistivities (resistances per unit distance) of the undifferentiated ground tissue and the already formed strands. If the resistivity of the ground tissue is high compared to that of the strands, the patterning process will produce the shortest connection between the incipient primordium and the existing vasculature, as the simple distance-minimizing algorithm would. However, if the discrepancy between the resistivities of the ground tissue and newly patterned strands is smaller and both decrease over time, the connections form according to the least resistance criterion and the threads align with a single family of parastichies, for appropriately chosen parameter values (Fig. S11b). An opposite set of parastichies can then be generated by applying, after some delay, the same resistance-minimizing algorithm, but excluding connections running close to those already formed (Fig. S11c). The resulting process reproduces the development and final structure of the reticulate vascular network characteristic of Bellis heads (compare Fig. 6 to Figs. 5g-i and 3g; see also Movies S2 and S3).

**Fig 6.**
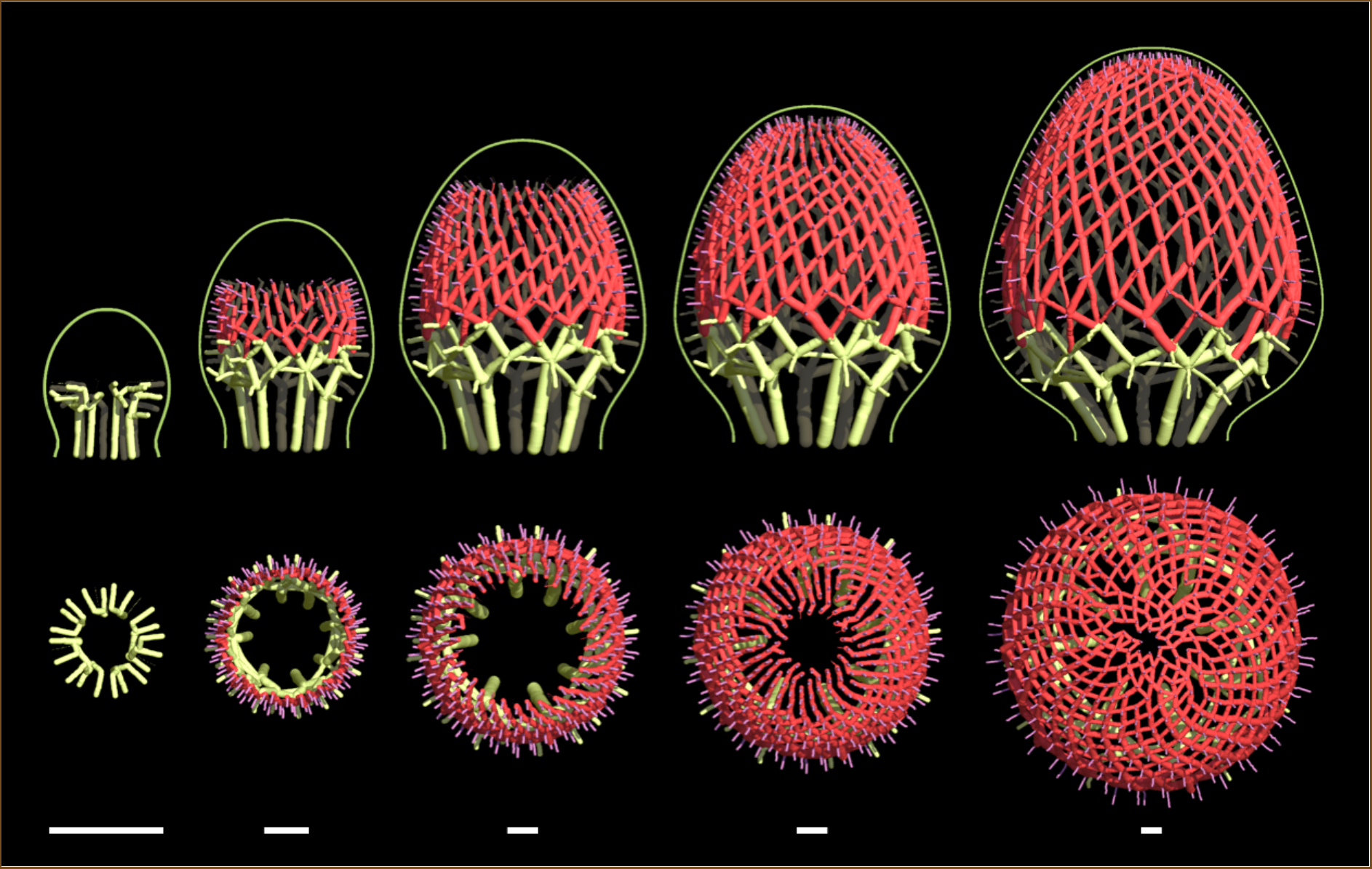
Simulated development of the Bellis head vasculature. Colors indicate abaxial strands (yellow-green), adaxial strands (red), and floret veins (purple). Scale bar: unit length.

Even in the highly regular vascular pattern of Bellis, some connections needed for “perfect” reticulation are missing (Fig. 3h). This phenomenon suggests that the secondary connections are formed with some probabilities, rather than deterministically. Moreover, in previous conceptual and computational models of vascular patterning it has often been postulated that the ground tissue may be polarized, biasing the formation of vascular strands in certain directions (e.g. Sachs, 1991, Bayer *et al*., 2009). In terms of our model, such bias can be captured by assuming that the resistivity of the ground tissue depends on directions, i.e., it is anisotropic. The theoretical morphospace (McGhee, 1999) of structures generated with different probabilities of secondary connections and different degrees of anisotropy on the Bellis head template (Fig. S12) includes patterns approximating the vasculature of the coneflower (Figs. 4f and S4c), Tanacetum (Figs. 4i and S4d), Craspedia (Fig. 4l and S4e), and Echinops (Fig. 4o and S4f) heads.

Depending on the head geometry and resistivity values, the paths of least resistance need not run close to the adaxial head surface, but may penetrate the ground tissue more deeply, connecting directly to the abaxial strands (Fig. S13a-d). Such paths approximate the vasculature observed in the sunflower (Figs. 3a-d and 5e,f) and thistle (Fig. 4c) heads. To model the observed irregular paths of individual strands in the sunflower (Fig. 7 and Movies S4-S6) we assumed that the direction in which subsequent segments are added is subject to significant random variation. In addition to the irregularities, the random variation of path extensions promotes the merging of adjacent paths, yielding the highly branched vascular patterns observed in real heads (Fig. S13e,f).

**Fig. 7.**
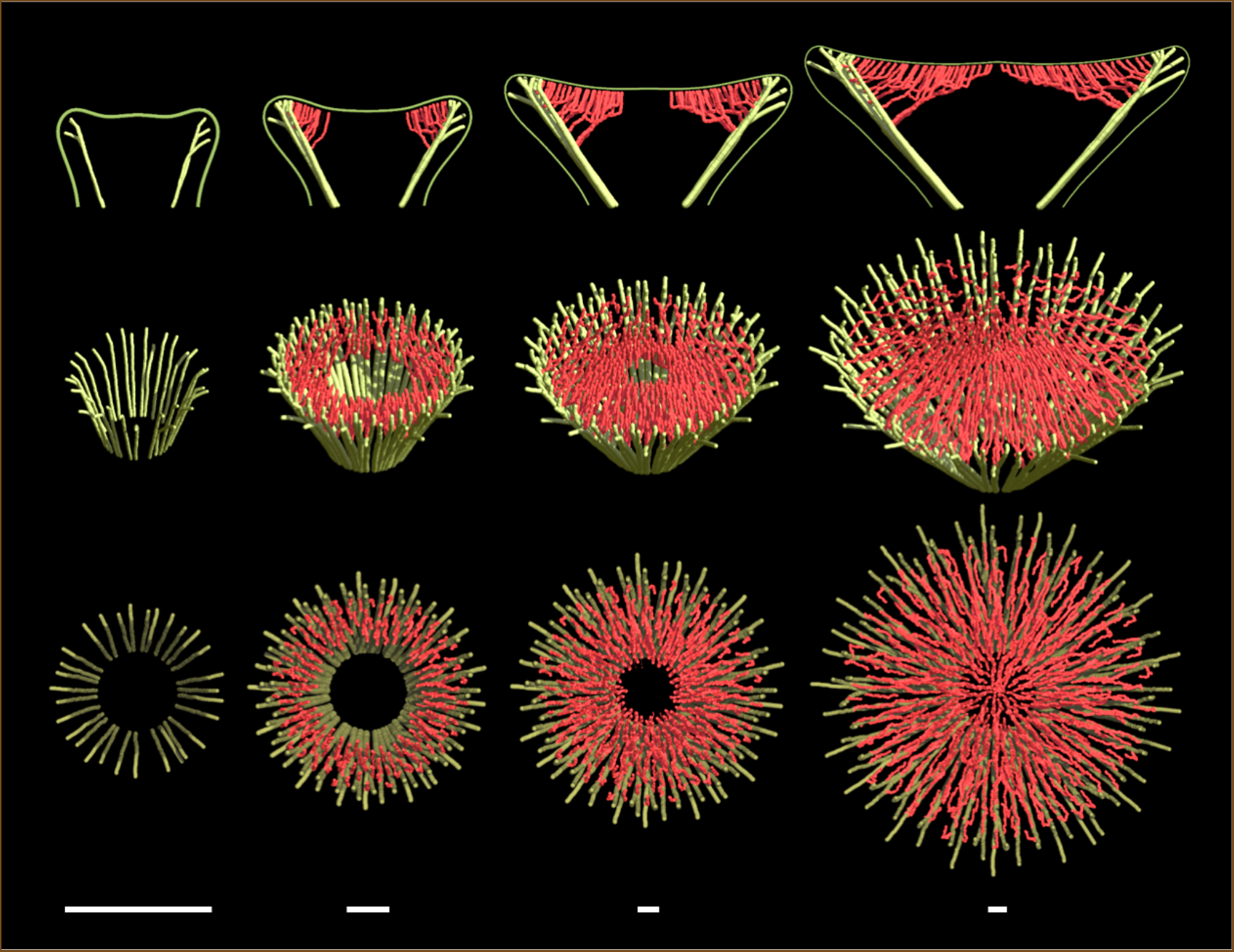
Simulated development of the sunflower head vasculature. Colors distinguish abaxial strands (yellow-green) from those originating in the florets (red). Scale bar: unit length.

In the cases discussed so far, the development of the vascular system was driven by the emergence of primordia. However, the polarization of ground tissue introduces a reference direction, in which adaxial strands may form ahead of the florets and thus independently of them. The underlying biological mechanism may involve a diffuse region of elevated auxin concentration similar to that observed in the shoot apical meristems of Arabidopsis (Barbier de Reuille *et al*., 2006; Galvan-Ampudia *et al*., 2020) towards which the adaxial strands extend. The minimum resistance algorithm then connects floret veins to the nearby adaxial strands, irrespective of the phyllotactic pattern. The resulting vasculature is characteristic of gerbera (compare Figs. 8 and 5a-c; see also Movies S7-S9).

**Fig. 8.**
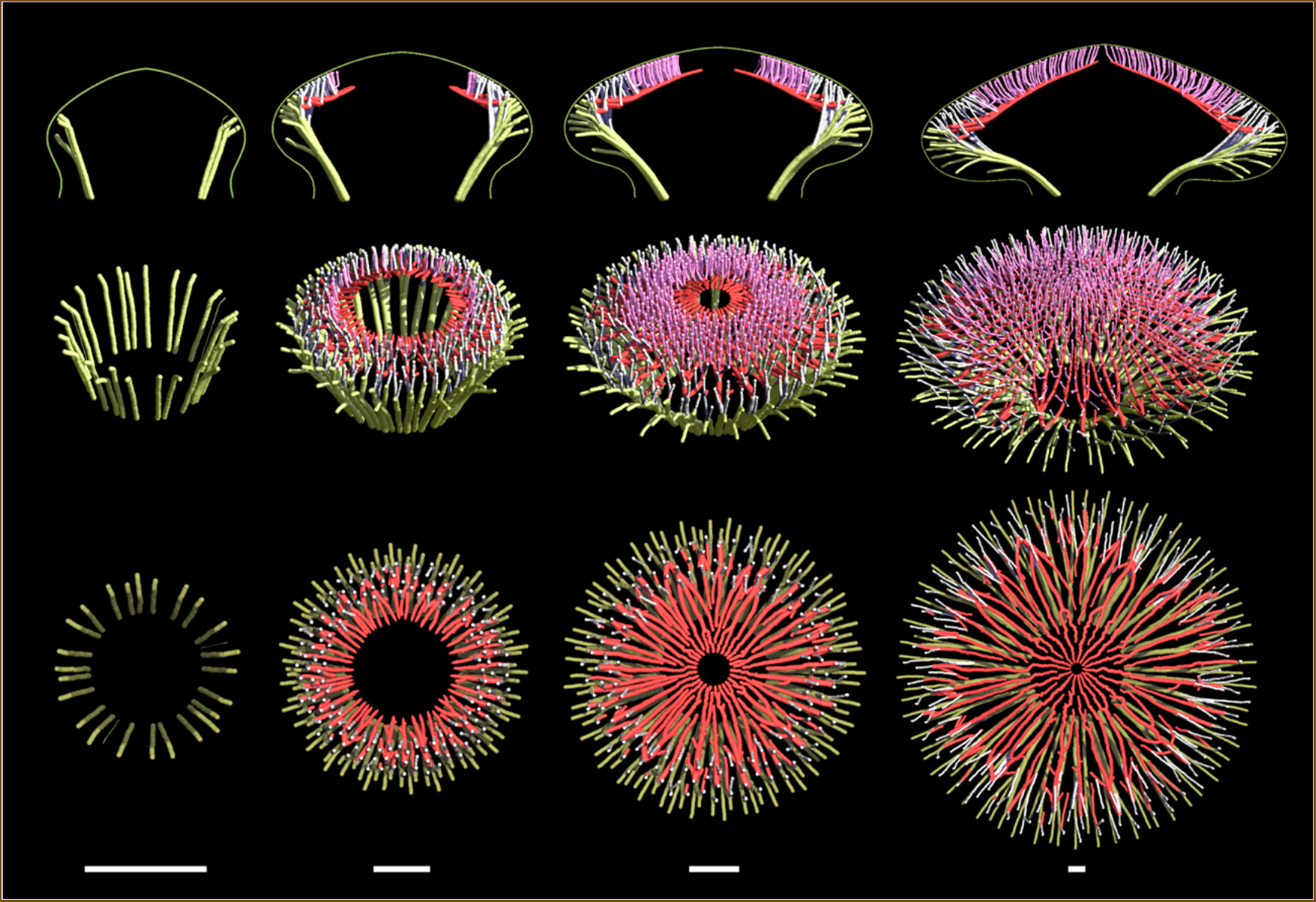
Simulated development of the gerbera head vasculature. Colors distinguish abaxial strands (yellow-green), adaxial strands (red), and floret veins (purple). Disk florets and floret veins have been suppressed in the bottom row to expose adaxial strands. Scale bar: unit length.

## Discussion

We have applied laboratory- and synchrotron-radiation-based micro-CT to explore vascular systems of flower heads. The throughput and resolution of micro-CT as well as the ease of sample preparation open the door to the effective screening of vascular structures in plants at different developmental stages. To analyze and annotate (segment) the complex vascular structures in heads, we developed two interactive programs. SHVR (Gu, 2022) augments images on a standard two-dimensional monitor with tactile feedback, giving the user the impression of touching individual vascular strands. ViNE (Hart, 2020) applies virtual reality technology to view and analyze structures directly in 3D.

Our results are summarized in Fig. 9. Using SR-μCT, we have confirmed previous observations of two distinct types of vascular patterns in heads: the reticulate pattern aligned with parastichies exemplified by Bellis (Philipson, 1946); and the sectorial pattern found in sunflower (Durrieu *et al*., 1985). In the latter case, strands originating in nearby florets form sympodial branching structures that deeply penetrate the ground tissue and connect directly to the abaxial strands. A similar pattern is characteristic of thistle heads. Nevertheless, most observed vascular patterns were shallow, running close to the receptacle surface. Among the heads we have examined, the coneflower was the clearest shallow counterpart of the sunflower, with the sympodial branching structures originating in the florets and connecting to the abaxial veins near the head perimeter. Craspedia, Echinops and Tanacetum heads combined features of shallow sectorial and reticulate patterns: the course of vascular strands was biased sectorially to some extent, while the patterns included loops, although not as regular as in Bellis. A relative outlier was the sectorial pattern of gerbera, characterized by the presence of radially oriented adaxial strands, to which floret veins connect in a monopodial branching pattern.

**Fig. 9.**
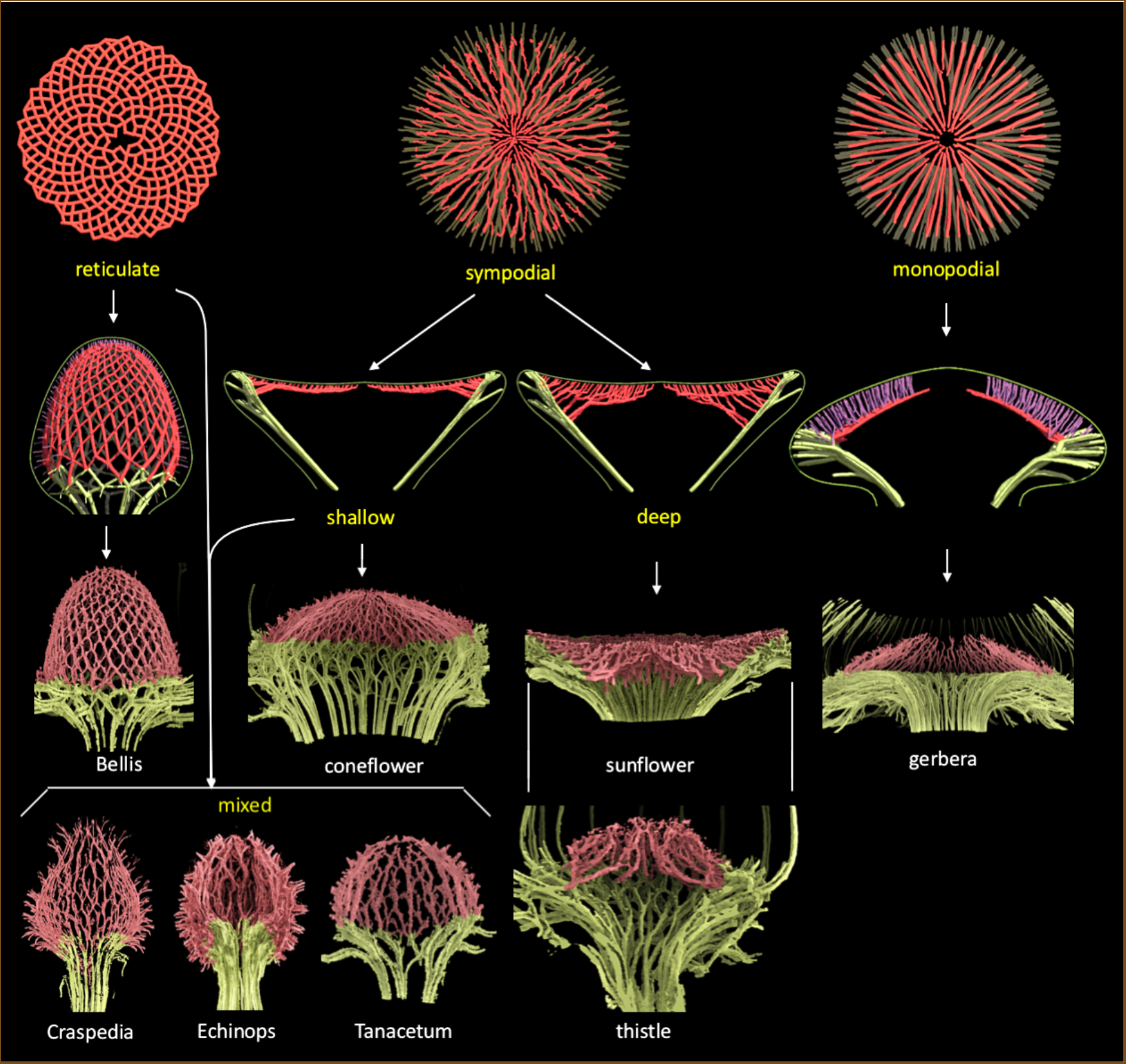
Relations between vascular structures in the analyzed heads.

To obtain insight into the developmental mechanisms that may produce the observed patterns, we constructed a dynamic model of vascular patterning in flower heads expressed in geometric terms (Owens *et al*., 2016; Cieslak *et al*., 2021). A key explicit assumption was that the vascular strands supporting new primordia extend the network formed so far in a manner minimizing the resistance of the resulting paths to the transport of auxin (Runions *et al*., 2017). To explain the vascular pattern in gerbera, we additionally assumed the presence of a region that attracts the emerging vascular strands, located in the central zone of the apex. With different, time-dependent resistivities of the vascular and ground tissues, our model reproduced the vascular patterns of all observed heads.

The eight sample species analyzed in this paper are but a minute fraction of the estimated 25,000–35,000 Asteraceae species (Mandel *et al*., 2019). This disparity leads to the question of whether there are further types of vascular patterns in heads fundamentally different from those encompassed by our study (Fig. 9). If the diversity of vascular patterns in heads echoes that of vein patterns in leaves, we should expect more patterns, even within the confines of our resistance-based model. For instance, our model can readily generate a non-reticulate pattern aligned with a single family of parastichies (Fig. S11b), or a pattern in which each floret connects directly to the base of the head though its own vascular strand (resulting in a vascular structure that resembles the branching architecture of an umbel inflorescence). The efficiency of micro-CT opens the door to addressing the question of vascular pattern diversity through a broad screening of heads.

The diversity of the vascular structures in heads raises the question of the molecular implementation of the patterning mechanism. According to current understanding, vascular patterns in plants result from feedback loops in which auxin regulates its own transport. In this process, maxima of auxin concentration emerge in the epidermis and initiate auxin flow into the subepidermal tissues. There, the flow canalizes into narrow paths that connect to the pre-existing vasculature, patterning new vascular strands (Jacobs, 1952; Sachs, 1969; Sachs, 1991; Reinhardt *et al*., 2003; Kang and Dengler, 2004; Scarpella *et al*., 2006; Wenzel *et al*., 2007; Bayer *et al*., 2009; O’Connor *et al*., 2014). The auxin reporter lines of gerbera indicate that auxin defines the sites of primordia in heads as it does in other model species (Zhang *et al*., 2021; Fig. S9), which suggests that the course of vascular strands in heads is patterned by a similar mechanism as well. Nevertheless, a more comprehensive analysis of the patterning of vasculature in heads at the molecular level is needed. The crucial next step will be the construction and analysis of transgenic Asteraceae plants reporting the spatio-temporal expression and cell-level distribution of auxin efflux carriers, the PIN1 proteins.

The prevalent view is that phyllotactic patterning of primordia takes place first, guiding the subsequent creation of vascular patterns. Nevertheless, an opposite possibility has been repetitively raised, suggesting that patterning information flows from the differentiated internal tissues towards the meristem surface and determines or influences organ positioning (Esau, 1954; Larson, 1975; Kang *et al*., 2003; Dengler, 2006; Banasiak and Zagórska-Marek, 2006; Kuhlemeier, 2007; Banasiak, 2011; Banasiak and Gola, 2023). It is difficult to imagine, however, how the development of highly regular phyllotactic patterns could be driven by locally irregular vascular strands. Likewise, it is not clear how the diversity of vascular patterns could lead to the uniformity of spiral phyllotactic patterns. In contrast, our models show that the observed vascular patterns, with large-scale organization but also with many local irregularities, can readily arise from regular phyllotactic patterns. Our observations and model thus support the prevalent view of the primacy of phyllotaxis over vascular patterning, although we cannot preclude a causality-blurring feedback loop, in which primordia and vascular strands are patterned concurrently.

The relation between the arrangement of florets and the underlying vascular pattern bears upon the long-standing question of the evolutionary origin of flower heads. Based on morphological studies of basal relatives of Asteraceae (Menyanthaceae, Goodeniaceae, and Calyceraceae), Pozner *et al*. (2012) suggested that the head-like structures in Calyceraceae and Asteraceae have evolved from branched inflorescences by the gradual reduction to a compact raceme. Considering the vascular system as a vestige of an ancestral free-standing branching system is consistent with our results. Gerbera, which appeared early in the evolution of Asteraceae (Bremer, 1994; Mandel *et al*., 2019), has heads with adaxial strands to which floret veins attach in a monopodial fashion characteristic of racemes. The patterns we observed in other, evolutionary later species could result from the reduction of these adaxial veins, causing a transition from monopodial to sympodial architecture. This switch could further lead to the alignment of vascular strands with parastichies, following a quantitative reduction of the sectorial bias. We are not certain, however, how much insight the branching architecture of presumed ancestral inflorescences has to offer. As previous work (Harder and Prusinkiewicz, 2013) and our results here show, both architectures may significantly vary without fundamentally affecting the arrangement of the flowers they support. Architectural transformations may therefore be incidental to the central question of how the shoot apical meristem evolved into a receptacle large enough to accommodate hundreds of florets arranged into stereotypical spiral phyllotactic patterns.

## Supporting information

Movie S1 - Gerbera fly through

Movie S2 - Bellis side

Movie S3 - Bellis top

Movie S4 - Sunflower side

Movie S5 - Sunflower high angle

Movie S6 - Sunflower top

Movie S7 - Gerbera side

Movie S8 - Gerbera high angle

Movie S9 - Gerbera top

## Acknowledgements

We thank Dr. Heikki Suhonen from the Department of Physics, University of Helsinki, for assistance with laboratory-based micro-CT, Chithra Kanurakaran for the suggestion of using SR-μCT for analyzing vascular patterns in plants and early guidance, and the staff at the Biomedical Imaging and Therapy facility at the Canadian Light Source^1^ for assistance. We also thank Sonny Chan for the introduction to haptics and help initiating the SHVR project, Usman Alim for sharing his expertise in volumetric rendering and segmentation, and Lynn Mercer for her work on the supplementary model description. This work was supported by the Plant Phenotyping and Imaging Research Centre/Canada First Research Excellence Fund (P.P. and M.C.), Academy of Finland Grants 1310318 and 1341774 (P.E.), and Natural Sciences and Engineering Research Council of Canada Discovery Grants 2014-05325 and 2019-06279 (P.P.). We also gratefully acknowledge the support of NVIDIA Corporation with the donation of two GPUs used in this research.

## Author contributions

PE, PP, TZ, and AO designed the research. TZ carried out experimental work and prepared all samples. TZ, JS, MC, PG, AO, PE, PP scanned samples. TZ, MC and JS reconstructed the data stacks. AO, PG and JH wrote the visualization software. TZ, PG, JH, AO, and MC visualized the data. AO and MC developed the models. PP, PE, AO, TZ, MC wrote the paper with input from coauthors. MC and TZ curated the data.

## Data availability

The reconstructed image stacks, segmentation masks, and polygon meshes reported in this paper are openly available from the University of Calgary’s OneDrive cloud storage (https://uofc-my.sharepoint.com). The data (150GB in total) are organized according to the order of appearance in the figures, as presented in Table S1. Image stacks are saved as multi-page TIF files, with segmentation information in MASK files specific to SHVR. Polygon meshes are in the PLY format, with segmentation information in CLR files specific to ViNE. Requests to access the shared drive should be made to the corresponding authors.

## Footnotes

1 The Canadian Light Source, a national research facility of the University of Saskatchewan, is supported by the Canada Foundation for Innovation (CFI), the Natural Sciences and Engineering Research Council (NSERC), the National Research Council (NRC), the Canadian Institutes of Health Research (CIHR), the Government of Saskatchewan, and the University of Saskatchewan.

