## Supplemental Information for "The Hidden Diversity of Vascular Patterns in Flower Heads"

The following Supporting Information is available for this article:

Supplemental Information (this file) including:
- Supplemental Figures S1-13
- Supplemental Table S1: Tomography, imaging techniques and data sets
- Supplemental Text S1: Details of synchrotron scanning

Supplemental Movies S1-S9
- S1: Fly-through of a scanned and segmented gebera head
- S2,S3: Simulated development of a Bellis head vasculature - side and top views
- S4-S6: Simulated development of a sunflower head vasculature - side, high-angle and top views
- S7-S9: simulated development of a gerbera head vasculature - side, high-angle and top views

**Supplemental Figures**

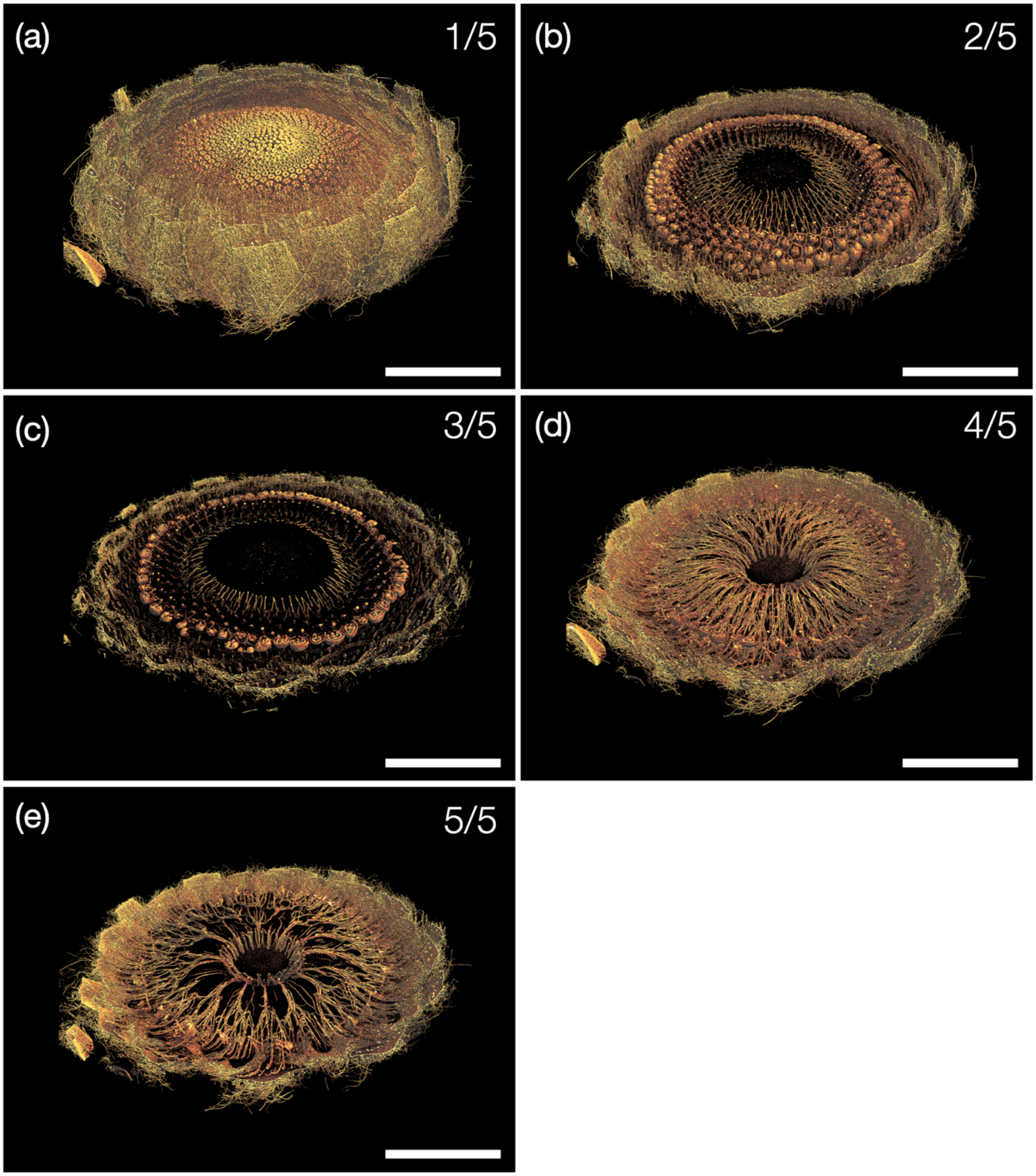


**Fig. S1** Volume rendering of a gerbera flower head. The fully developed head (Fig. 1a) was virtually sliced into horizontal sections showing the sectorial pattern of adaxial (b,c) and abaxial (d,e) vascular strands. Scale bars: 5 mm.

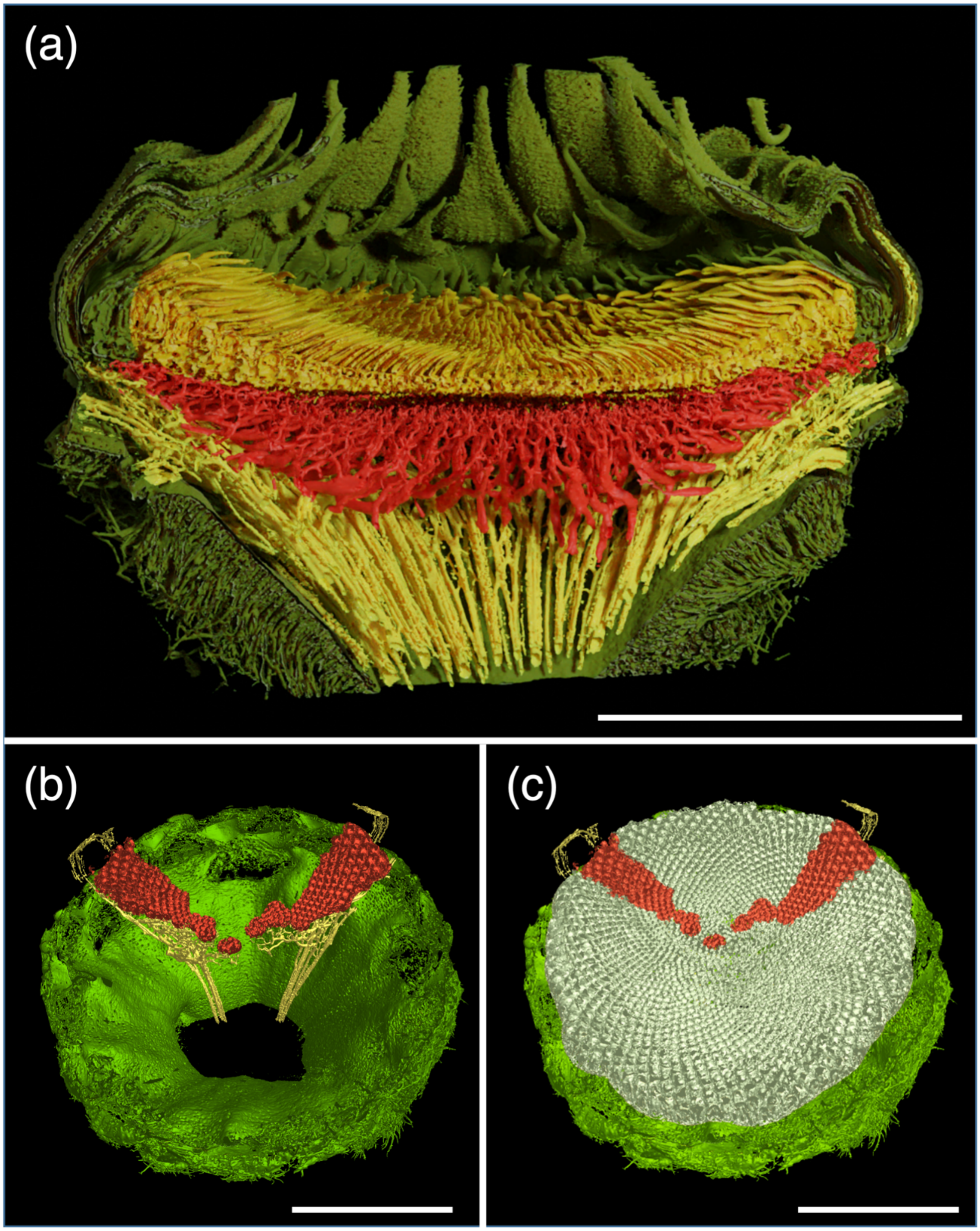


**Fig. S2** Relation between primordia and vasculature in a sunflower head. (a) Volumetric rendering of half of the head showing adaxial strands (yellow), and strands originating in the florets (red). (b) Two sample sectors of the vasculature (yellow) and the floret primordia (red) supported by them with the remaining florets digitally removed. (c) The same flower head as (d) with all florets displayed. Scale bars: 1 cm.

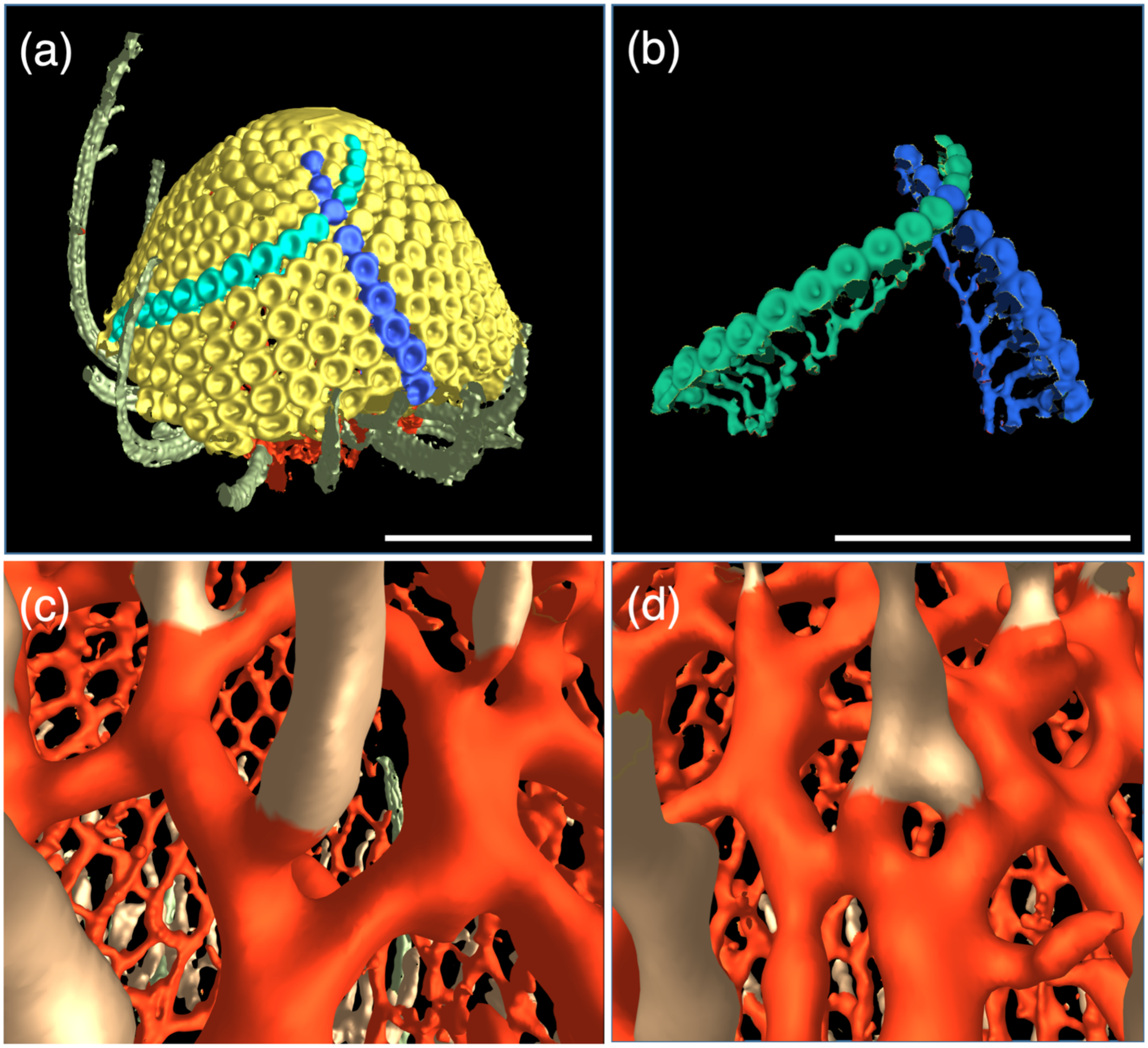


**Fig. S3** Relation between primordia and vasculature in a Bellis flower head. (a) External view of the head, with primordia forming two intersecting parastichies segmented in cyan and blue. (b) The same head with all but the two highlighted parastichies hidden. The underlying vasculature supports the floret primordia along these two parastichies. (c,d) Magnified views of the vascular system, showing that floret veins (light brown) may attach to the reticulate vasculature outside a junction between two strands running in the opposite directions (c), or connect to two strands (d)Scale bars (a, b): 1 cm, (c,d): variable scale due to perspective projection.

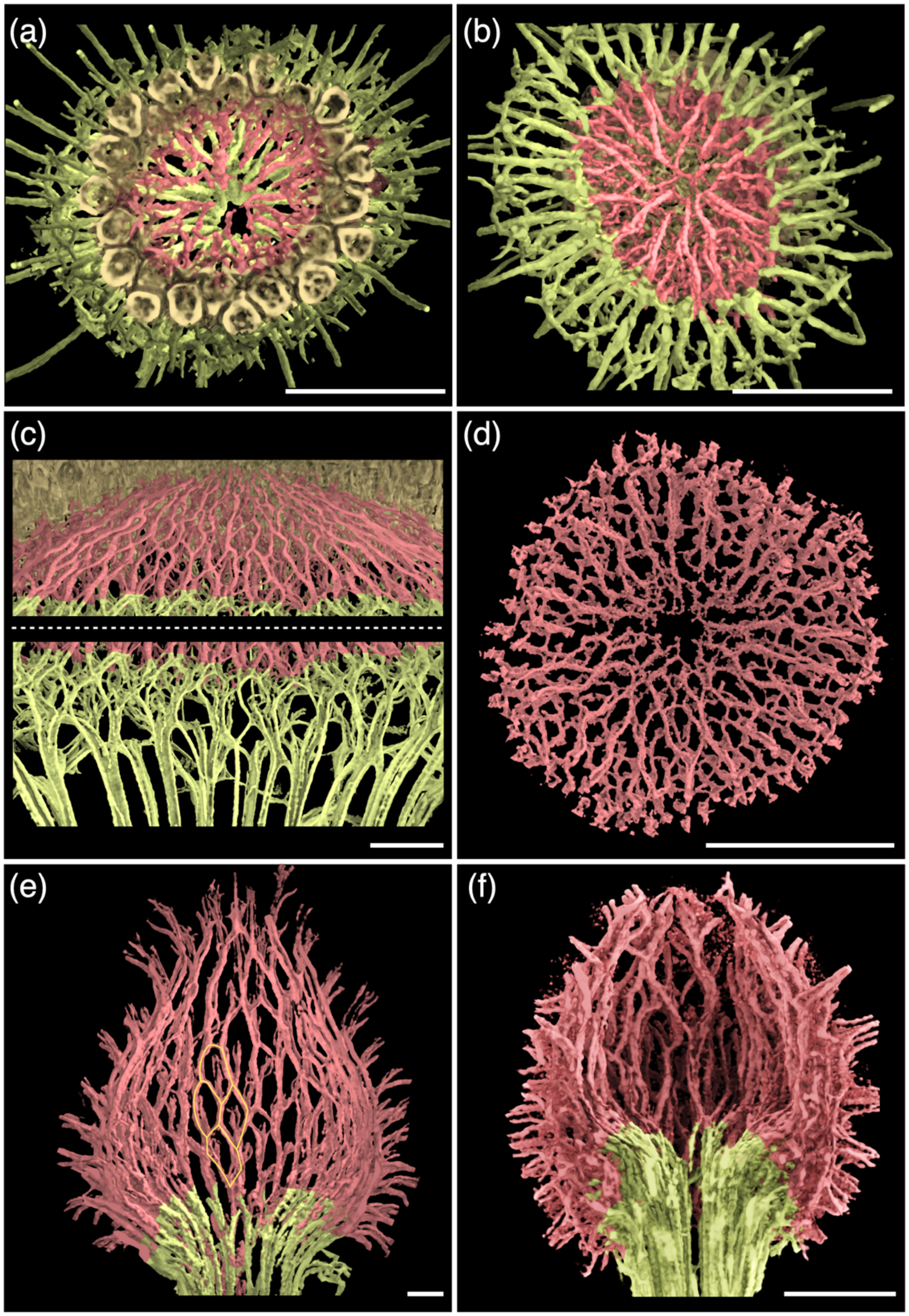


**Fig. S4.** Intermediate vascular system in flower heads shown in Fig 4. (a,b) Top and bottom view of vasculature strands in a thistle flower head showing sectorial arrangement of the vascular system. (c) Close-up views of floret strands (top) and abaxial strands (bottom) in a flower head of coneflower. (d) Veins of Tanacetum head showing an irregularly connected reticulate pattern. (e,f) Vascular systems in globular heads of Craspedia (e) and Echinops (f). A portion of the reticulate pattern in Craspedia is highlighted by yellow lines. Note the multiple floret veins entering each secondary head. Scale bar: 0.5 mm.

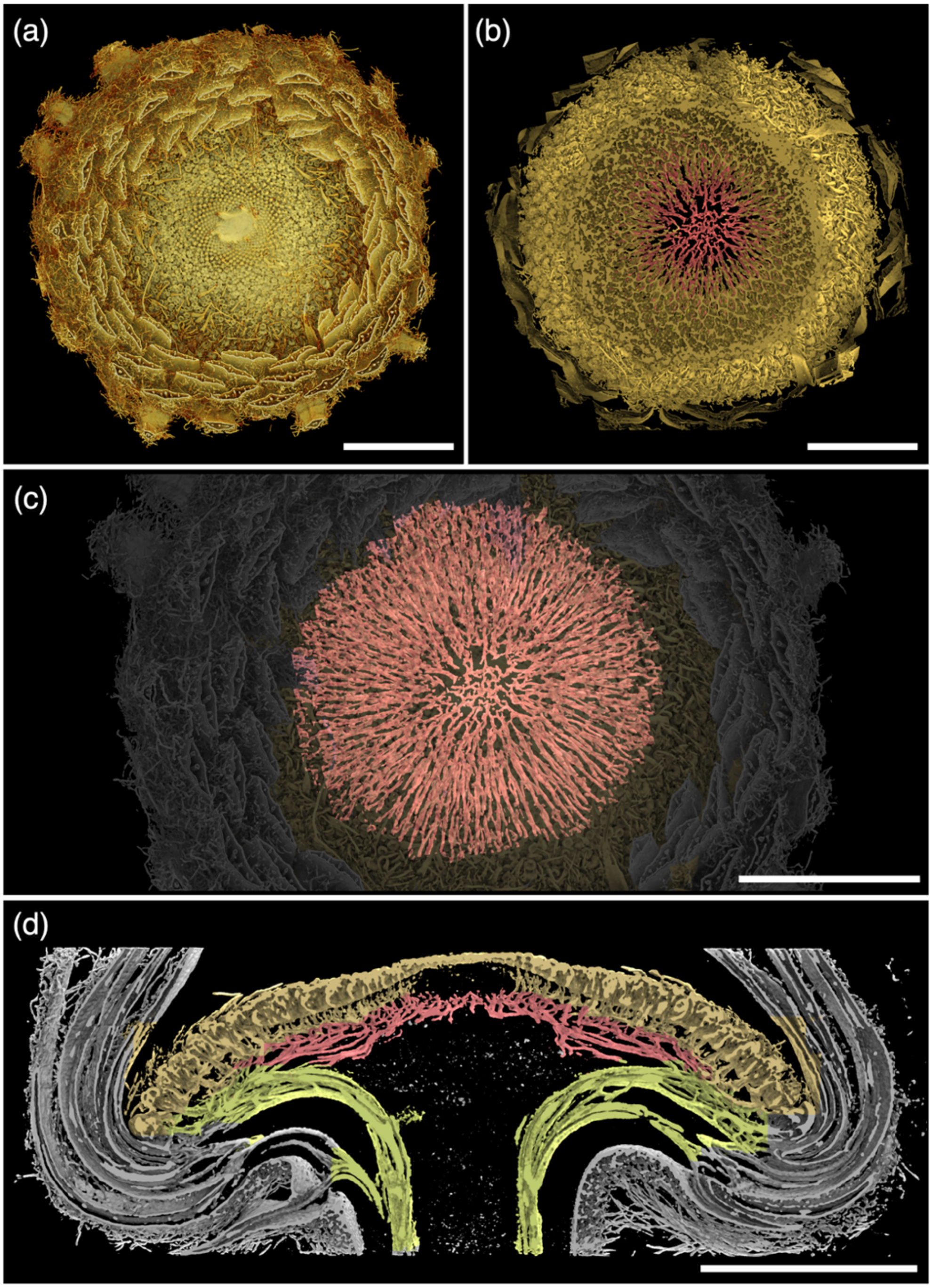


**Fig. S5** Vascular system in a transgenic gerbera head with a downregulated *SEPALLATA*-like *GRCD* gene. (a) An outside view showing the undifferentiated head centre. (b) A virtual section of the head in the same position, demonstrating the presence of the vasculature under the undifferentiated head center. (c) The complete adaxial vasculature superimposed on the head image at the same scale. (d) A thin longitudinal section demonstrating the presence of adaxial strands under the undifferentiated head center. The abaxial and adaxial vascular strands are colored green and red, respectively. The strands that penetrate bracts have not been segmented. Scale bars: 5 mm.

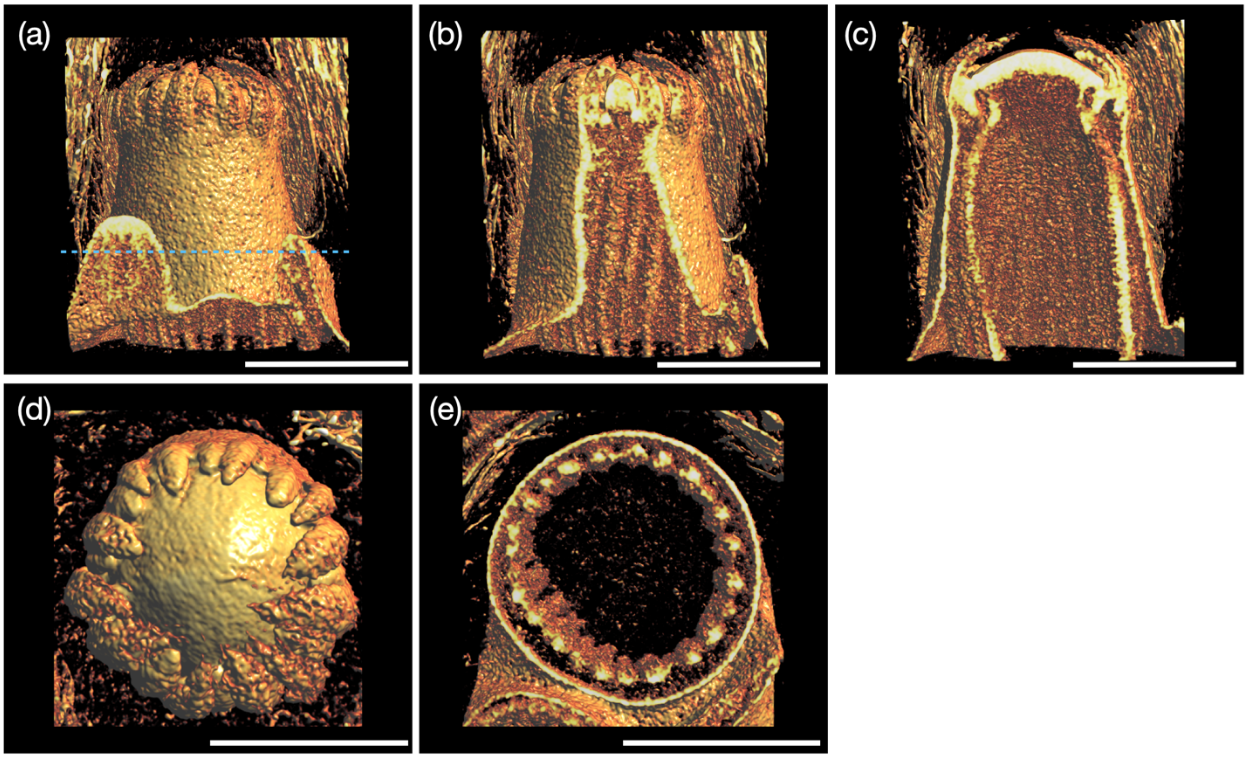


**Fig. S6** Volumetric rendering of a gerbera head meristem at an early stage of development. (a-c) Side view and two longitudinal sections with different positions of the cutting plane. (d,e) Top view and a horizontal cross-section showing the arrangement of vascular strands into a cambium ring. The cutting plane in (e) is marked with a blue dashed line in (a). Scale bar: 1 mm.

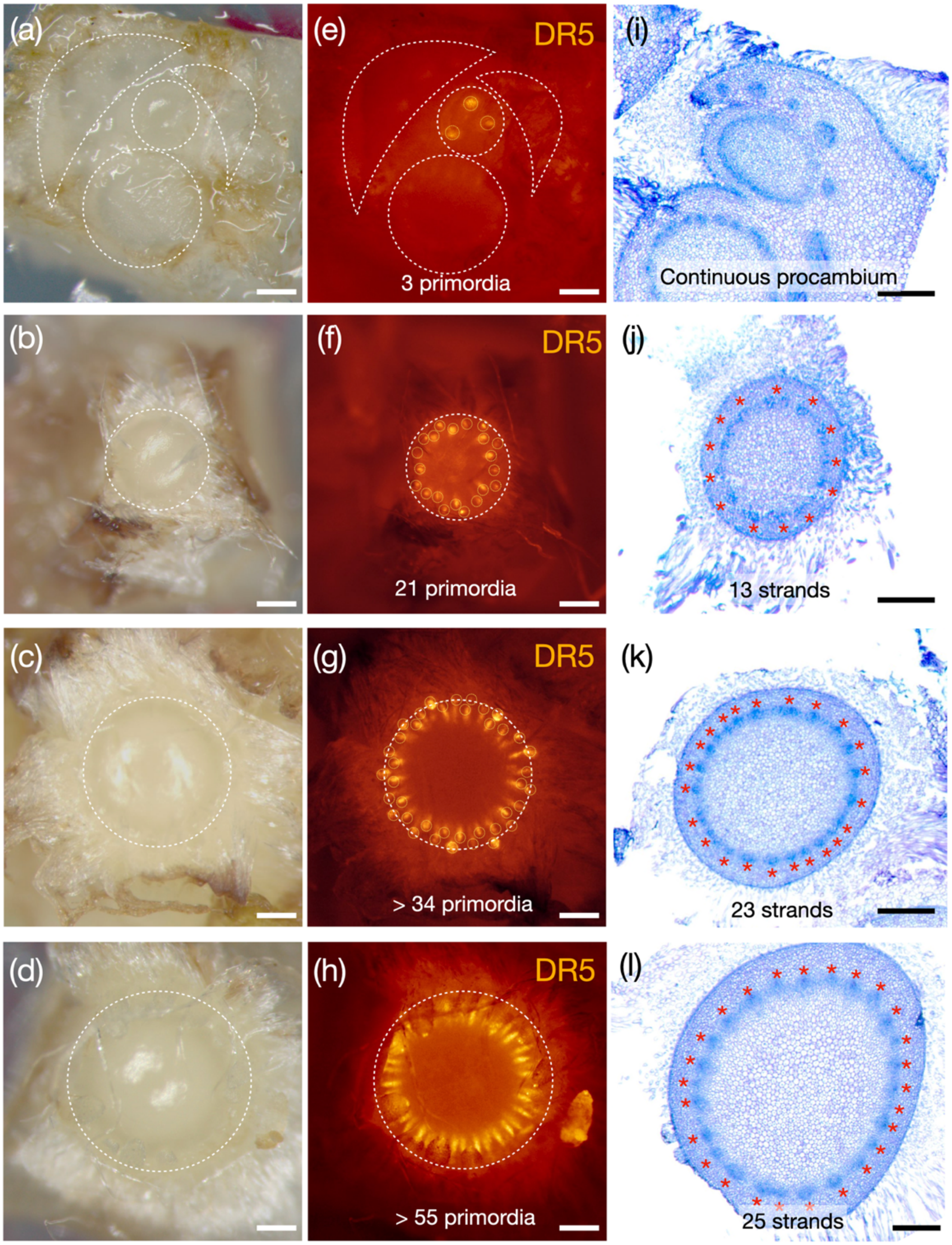


**Fig. S7** Association of auxin patterning with patterning of the vasculature. Four independent head meristem samples (a-d) from transgenic gerbera lines expressing the *DR5rev:3xVENUS-N7* auxin reporter (Zhang et al. 2021) show increasing numbers of DR5 maxima on the meristem surface corresponding to the emerging bract and floret primordia (e-h). The rightmost column (i-j) shows a representative histological, toluidine blue stained section underneath the surface of the same meristem, with vascular bundles counted and marked in asterisks. Scale bar: 200 µm.

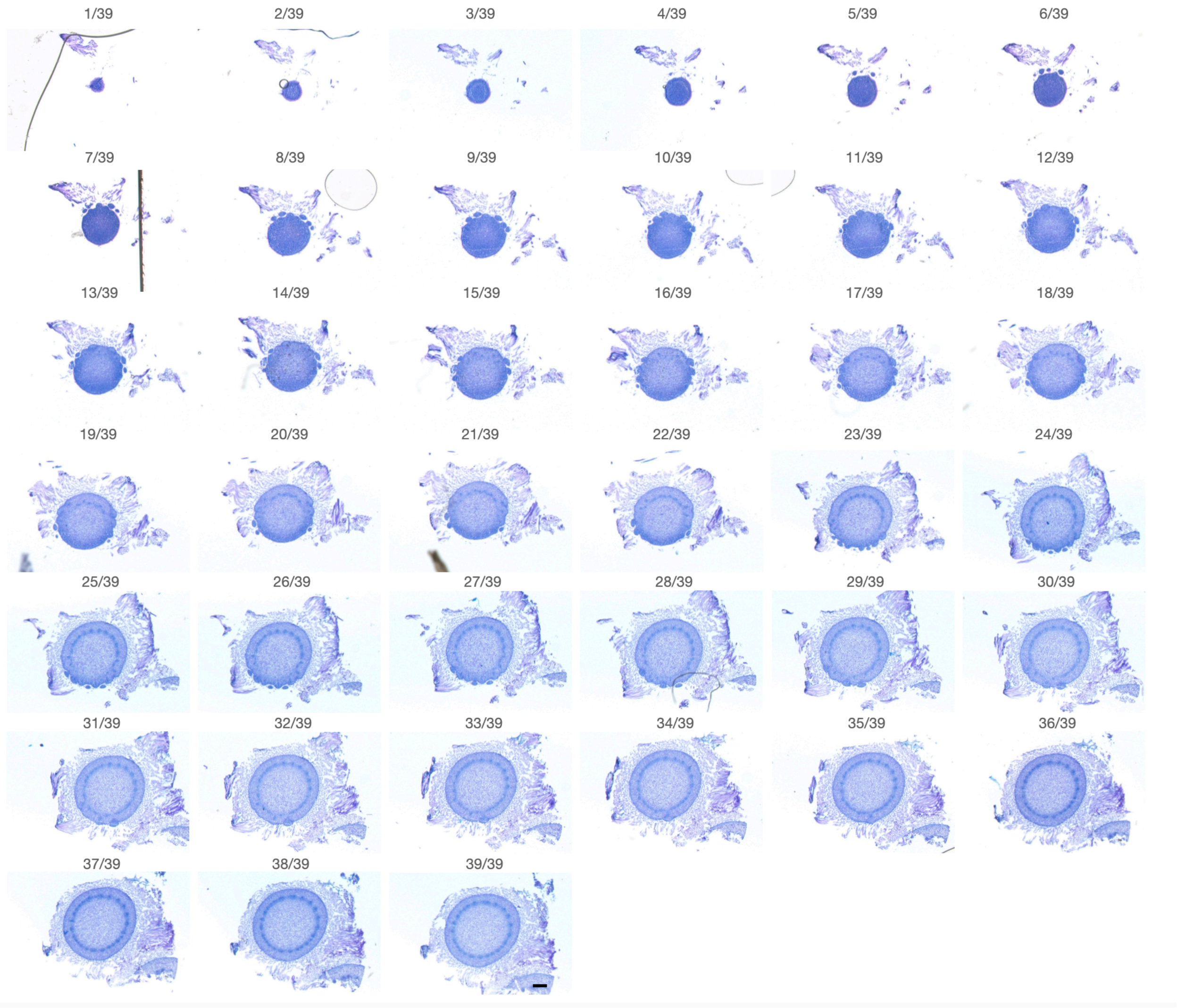


**Fig. S8** An example of serial section for histological analysis. The data represents the 39 slides collected from the sample corresponding to the Fig. S7c. Scale bar: 200 µm.

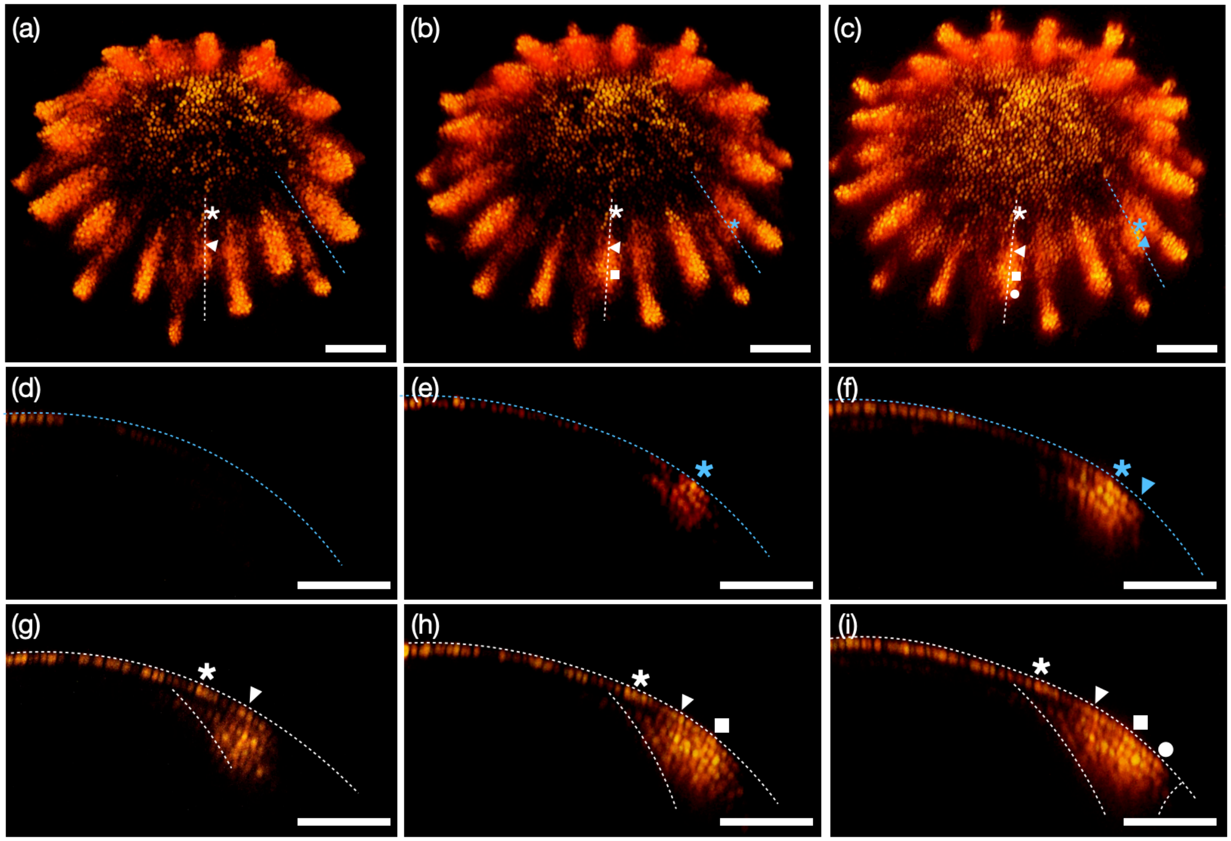


**Fig. S9** Auxin distribution in emerging primordia observed in transgenic gerbera lines expressing the DR5rev:3xVENUS-N7 reporter. (a-c) Live-imaging data of a gerbera head meristem from Zhang et al. (2021). Two primordia (marked by dashed white and blue lines) at 0h (a), 48h (b) and 92h (c) timepoints were selected to perform optical longitudinal sections. (d-f) Longitudinal optical cross-section of an emerging, early-stage primordium marked in blue in (a-c)). (g-i) Longitudinal optical cross-section of the relatively later stage primordium marked in white in (a-c). Corresponding cell positions are marked by asterisks, triangles, squares and circles. The auxin signal initiating primordia at the meristem surface propagates into subepidermal tissues as In meristems of plants not forming heads. Scale bar: 100 µm.

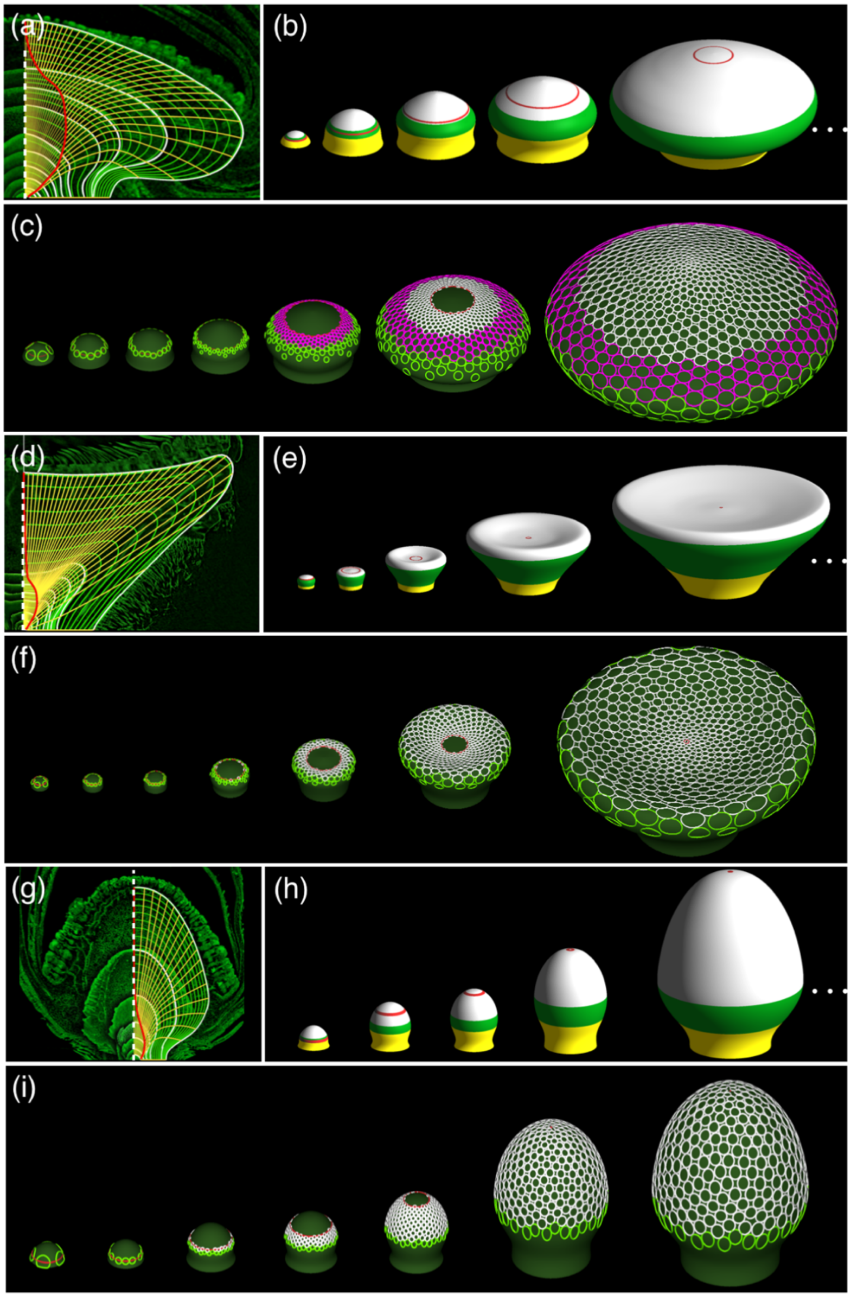


**Fig. S10** Construction of the head templates. (a,d,g) Superimposed virtual longitudinal sections of scanned gerbera (a), sunflower (d) and Bellis (g) heads (green) in different stages of development are manually traced to obtain receptacle contours (white), trajectories of points on the receptacle surface (yellow) and position of the active ring (red) as the heads grow. (b,e,h) Snapshots of the resulting data-driven simulations of the head development. Colors indicate the adaxial (white) and abaxial (green) portion of the receptacle surface, the top portion of the supporting stem (yellow), and the position of the active ring (red). (c,f,i) Snapshots of head development with simulated phyllotactic patterns. Pink in (c) indicates trans florets. Volumetric representations of these heads provide the templates for simulating vascular pattern development.

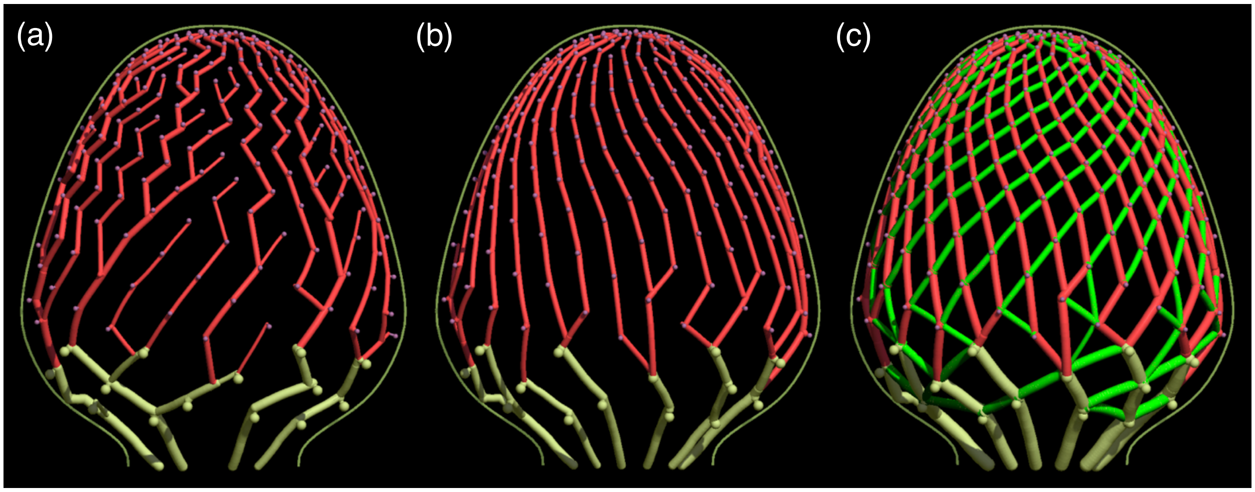


**Fig. S11** The key conceptual steps leading to the model of vascular patterning in Bellis heads. (a) Distance-based model produces an irregular branching pattern of strands that switch between parastichies. (b) Resistance-based model produces an open pattern with vascular strands consistently following one family of parastichies. (c) Additional connections (green) result in the regular reticulate pattern characteristic of Bellis.

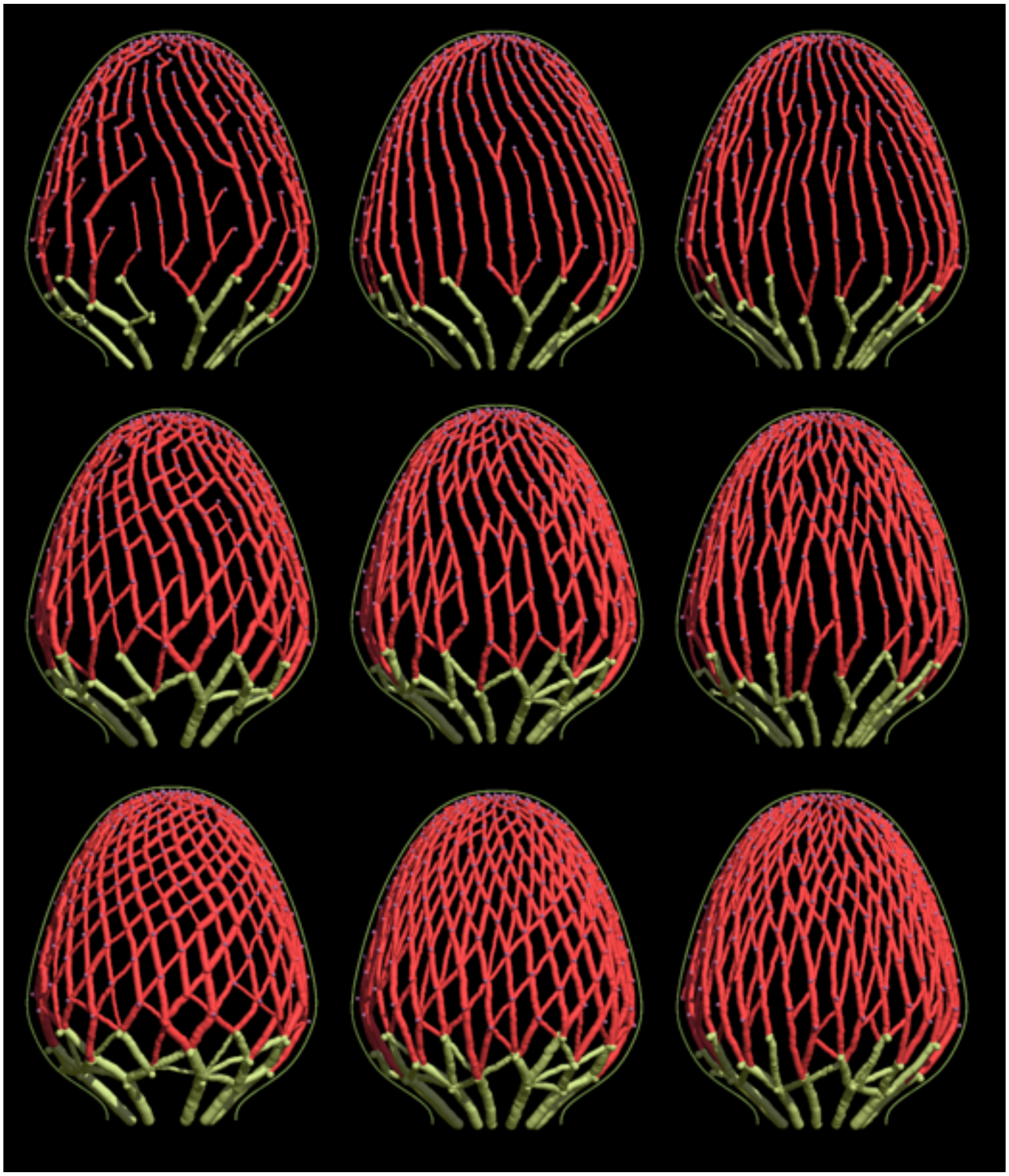

**Fig. S12** The impact of increasing sectoriality (left to right) and reticulation (top to bottom) on vascular patterns simulated using the Bellis head template. All remaining parameters have been kept constant.

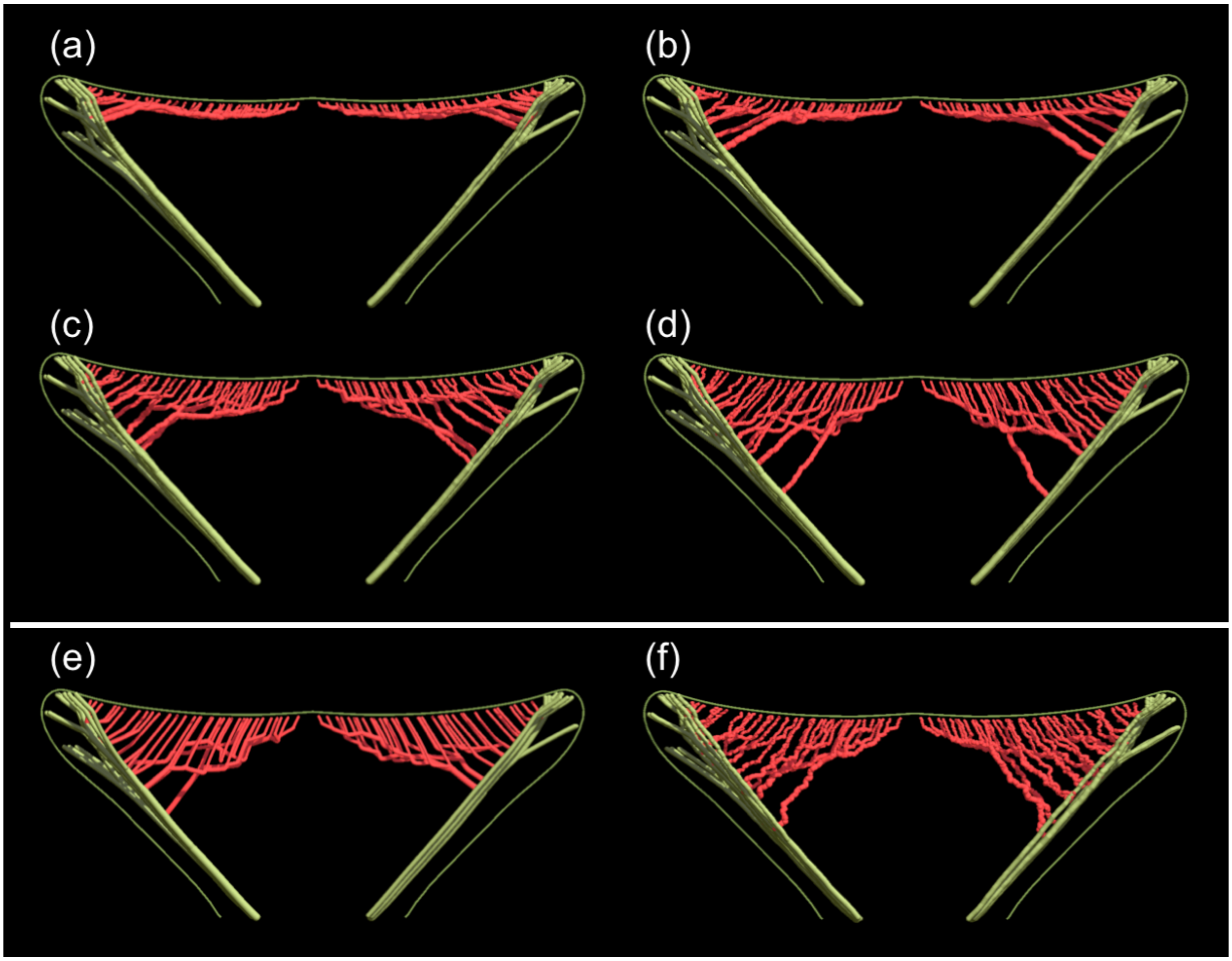


**Fig. S13** Exploration of the parameter space of sympodial vasculature simulated on the sunflower head template. (a-d) Decreasing resistance of the ground tissue to the auxin transport results in increasingly deep penetration of the ground tissue. (e,f) Increased random changes in the path direction lead to a more branched pattern.

**Table S1** Tomography, imaging techniques and data sets used to visualize scanned heads in this paper.

| Figure | | Method | Res. | Data | Visualization method |
|---|---|---|---|---|---|
| **Fig 1** | b,c,d | SR-μCT | 13 μm | Gerbera1.tif<br>Gerbera1.mask | SHVR (b,d)<br>SHVR segmented (c) |
| **Fig 2** | a-e | SR-μCT | 13 μm | Gerbera2.ply<br>Gerbera2.clr | ViNE segmented + Blender (a,b,c)<br>ViNE segmented (d,e) |
| **Fig 3** | b,c | SR-μCT | 13 μm | Sunflower1.tif<br>Sunflower1.mask | SHVR (b)<br>SHVR segmented (c) |
| | d | SR-μCT | 13 μm | Sunflower1.ply<br>Sunflower1.clr | ViNE segmented + Blender |
| | f,g | SR-μCT | 3.61 μm | Bellis1.tif<br>Bellis1.mask | SHVR (f)<br>SHVR segmented (g) |
| | h | SR-μCT | 3.61 μm | Bellis1.ply<br>Bellis1.clr | ViNE segmented + Blender |
| **Fig 4** | b,c | SR-μCT | 3.61 μm | Thistle.tif<br>Thistle.mask | SHVR (b)<br>SHVR segmented (c) |
| | e,f | SR-μCT | 3.61 μm | Coneflower.tif<br>Coneflower.mask | SHVR (e)<br>SHVR segmented (f) |
| | h,i | SR-μCT | 3.61 μm | Tanacetum1.tif<br>Tanacetum1.mask | SHVR (h)<br>SHVR segmented (i) |
| | k,l | SR-μCT | 13 μm | Craspedia.tif<br>Craspedia.mask | SHVR (k)<br>SHVR segmented (l) |
| | n,o | SR-μCT | 3.61 μm | Echinops.tif<br>Echinops.mask | SHVR (n)<br>SHVR segmented (o) |
| **Fig 5** | a | Skyscan 1272 | 5 μm | GerberaDev1.tif<br>GerberaDev1.mask | SHVR segmented |
| | b | GE Phoenix Nanotom | 7 μm | GerberaDev2.tif<br>GerberaDev2.mask | SHVR segmented |
| | c | GE Phoenix Nanotom | 10 μm | GerberaDev3.tif<br>GerberaDev3.mask | SHVR segmented |
| | d | Skyscan 1272 | 5 μm | SunflowerDev1.tif<br>SunflowerDev1.mask | SHVR segmented |
| | e | Skyscan 1272 | 5 μm | SunflowerDev2.tif<br>SunflowerDev2.mask | SHVR segmented |
| | f | Skyscan 1272 | 5 μm | SunflowerDev3.tif<br>SunflowerDev3.mask | SHVR segmented |
| | g | SR-μCT | 3.61 μm | BellisDev1.tif | SHVR segmented |

| | | | | | |
|---|---|---|---|---|---|
| | | | | BellisDev1.mask | |
| | h | SR-μCT | 3.61 μm | BellisDev2.tif<br>BellisDev2.mask | SHVR segmented |
| | i | SR-μCT | 3.61 μm | BellisDev3.tif<br>BellisDev3.mask | SHVR segmented |
| **Fig S1** | a-e | SR-μCT | 3.61 μm | Gerbera1-3_61.tif | SHVR |
| **Fig S2** | a | SR-μCT | 13 μm | Sunflower1.ply<br>Sunflower1.clr<br>(Same as Fig. 3d) | ViNE segmented + Blender |
| | b,c | SR-μCT | 13 μm | Sunflower1.ply<br>Sunflower1S.clr | ViNE segmented |
| **Fig S3** | a-d | SR-μCT | 3.61 μm | Bellis1.ply<br>Bellis1S.clr | ViNE segmented |
| **Fig S4** | a,b | SR-μCT | 3.61 μm | Thistle.tif<br>Thistle.mask<br>(Same as Fig. 4b,c) | SHVR segmented |
| | c | SR-μCT | 3.61 μm | Coneflower.tif<br>Coneflower.mask<br>(Same as Fig. 4e,f) | SHVR segmented |
| | d | SR-μCT | 3.61 μm | Tanacetum2.tif<br>Tanacetum2.mask | SHVR segmented |
| | e | SR-μCT | 3.61 μm | Craspedia.tif<br>Craspedia.mask<br>(Same as Fig. 4k,l) | SHVR segmented |
| | f | SR-μCT | 3.61 μm | Echinops.tif<br>Echinops.mask<br>(Same as Fig. 4n,o) | SHVR segmented |
| **Fig S5** | a-d | SR-μCT | 13 μm | GRCD.tif<br>GRCD.mask | SHVR segmented |
| **Fig S6** | a-e | Skyscan 1272 | 5 μm | GerberaDev1.tif<br>(Same as Fig. 5a) | SHVR |

Note: Samples scanned using laboratory or SR μCT at 13 microns were reconstructed with NRecon, samples scanned using SR-μCT 3.61 microns were reconstructed with UFO-KIT. Polygonal mesh (.ply) files were obtained from the .tif files with the same names using the IsoPoly program. The .mask and .clr files contain segmentation information obtained using SHVR and ViNE, respectively.

**Supplemental text S1: Details of synchrotron scanning**

We used two different high-resolution X-ray imaging systems to scan flower heads at the Canadian Light Source Biomedical Imaging and Therapy SR-µCT facility.

The first system was M11427-62 by Hamamatsu Photonics K.K. (Hamamatsu City, Japan) including an ORCA-Flash4 Digital sCMOS camera with a 2048x2048 pixel array and a 6.5x6.5 $\mu m^2$ pixel size, coupled through an AA60 microscope with a x0.5 objective to a 200 µm thick cerium-doped lutetium aluminum garnet (LuAG:Ce) scintillator, which gave an effective pixel size of 13 µm and a field of view of 26.6 mm (horizontal) by 4 mm (vertical, limited by the beam divergence). The samples were glued to the top of a standard Falcon tube and placed ~10 cm from the camera.

The second system consisted of a pco.edge 5.5 sCMOS camera by PCO AG (Kelheim, Germany), with a 2560x2160 pixel array and 6.5x6.5 $\mu m^2$ pixel size, coupled through an Optique Peter (Lentilly, France) microscope with a x1.8 objective to the same scintillator as above, which gave an effective pixel size of 3.61 µm and a field of view of 9.24 mm (horizontal) by 4 mm (vertical). Each sample was mounted on a Goniometer Head 1005 by Huber Diffraktiontechnik GmbH & Co. KG (Rimsting, Germany) and placed 5 cm from the camera.

During the scanning, each sample was rotated from 0 to 180° in 0.06° increments around the sample's approximate axis of symmetry, producing 3000 projection images per 4 mm vertical section. Samples higher than 4 mm were scanned by combining 2 to 4 vertical sections. For samples that were too wide (diameter over 9 mm at pixel size of 3.61 µm), we used half-view acquisition mode, in which the samples were offset horizontally relative to the detector's field-of-view and rotated 360° around an axis near the sample's edge, yielding twice as many projection images (6000) per each 4 mm vertical section of the sample. Pairs of half-view projections lying in the same planes were then stitched together along the sample's axis of rotation, resulting in 3000 full projections with the horizontal field of view increased to approximately 18 mm. Finally, we collected 10 dark signal images (no X-ray beam on the detector) and 10 flat signal images (with the X-ray beam on but no sample) to allow for compensating imperfections in the imaging system and normalize the intensity of projected images.
